# Redesigning petal shape, size, and color in soybean reveals unexpected phenotypes for floral organ development

**DOI:** 10.1101/2025.06.05.657847

**Authors:** Nicole Szeluga, Noor AlBader, Samantha Pelletier, Kylie Weis, Arielle Johnson, Noah Fahlgren, Mikhaela Neequaye, Gus Vogt, Ryan DelPercio, Patricia Baldrich, Kelsey J.R.P. Byers, Blake C. Meyers, Margaret H. Frank

## Abstract

Soybean (*Glycine max*) has not yet benefited from large-scale hybrid breeding efforts due to its small, self-fertilizing flowers that are difficult to emasculate, and limited attractiveness to pollinators. This study explores targeted floral trait engineering to enhance pollinator attraction, aiming to overcome barriers to soybean hybridization. We generated a high-resolution floral organ expression atlas and H3K4 trimethylation ChIP-Seq dataset to identify candidate genes involved in petal development, nectar sugar content, and petal pigmentation. Using CRISPR-based activation and repression systems, we modified the expression of *AINTEGUMENTA (GmANT)*, *BIGPETAL (GmBPE)*, and *SUCROSE TRANSPORTER2 (GmSUC2)*. Contrary to expectations based on *Arabidopsis* homologs, transcriptional activation of *GmANT_B* and repression of *GmBPE* led to reduced, rather than increased, petal size, highlighting divergent regulatory mechanisms in soybean. Complementation of the *W1* gene that controls petal pigmentation successfully converted white petals to purple, with preliminary evidence indicating that this color conversion may increase pollinator visitation. These results underscore the complexity of floral development in soybean and provide foundational tools and resources for future efforts to engineer reproductive traits for hybrid seed production.

## Introduction

Hybrid breeding for heterosis has been used to boost yields in many crops, including maize, rice, and canola (Fu *et al.*, 2014; Labroo *et al.*, 2021; Li *et al.*, 2008; Sleper *et al.*, 2006). Despite its agricultural prominence, soybean (*Glycine max*) is grown as an inbred crop that has yet to benefit from heterosis (American Soybean Association; Dohlman and Hansen, 2022; National Agricultural Statistics Service, 2022). Hybrid breeding is particularly challenging in soybean because the plant produces small flowers that are typically self-fertilized prior to opening, and they are overall less attractive to pollinators needed for large-scale hybrid breeding. In our previous work, we addressed this first barrier using a biotechnology approach, introducing barnase/barstar, a rescuable male-sterility system that blocks self-fertilization in soybean (Szeluga *et al.*, 2023). Data from field studies show that bee pollinator-driven outcrossing does occur amongst soybean varieties, albeit at low rates, indicating that this second barrier to hybrid breeding could be addressed by enhancing floral traits that attract pollinators (Ahrent and Caviness, 1994). In this study, we address this second barrier to hybrid breeding, by testing whether a targeted redesign of soybean petal and reward features can be used to enhance pollinator recruitment.

Floral traits, including petal size and color, flower number, volatile profile, and nectar sugar content are known factors that influence pollinator attraction (Severson and Erickson, 1984). Bees, including honeybees, are the most likely pollinators for field-grown soybean and are known to prefer darker-colored petals, higher percent sugars in nectar (between 40-60%), and larger floral displays (Balfour and Ratnieks, 2023; Blettler *et al.*, 2018; Giurfa *et al.*, 1995; Leleji, 1973; Waller, 1972). Because honeybees have photoreceptors most sensitive to the UV/blue region of the light spectrum, they are more attracted to floral displays emitting these wavelengths (Giurfa *et al.*, 1995; Leleji, 1973). Soybeans generally produce either white or purple flowers, reflecting constrained genetic variation in the genes that control flower color. The *W1* gene (the soybean version of the anthocyanin biosynthesis pathway biosynthetic gene *F3’5’H*) is one of the six genes primarily responsible for producing anthocyanin pigmentation in soybean petals (Sundaramoorthy *et al.*, 2016, 2021; Yan *et al.*, 2014). Soybean varieties with white flowers, including the genetic model, Williams 82 (W82), carry a nonsense mutation in *W1*, resulting in a lack of anthocyanin production (Sundaramoorthy *et al.*, 2016, 2021). Similarly, petal size has been shown to influence pollinator visitation frequency. Larger flowers are more visible from longer distances, and bumblebees can forage faster from flowers with large floral displays (Moyroud and Glover, 2017).

Multiple studies in Arabidopsis have identified genes that show altered petal size when mutated, including *AINTEGUMENTA* (*ANT*), *BIGPETAL* (*BPE*), and *BIG BROTHER* (*BB*) (Brioudes *et al.*, 2009; Chen *et al.*, 2018; Disch *et al.*, 2006; Mizukami and Fischer, 2000; Randall *et al.*, 2015; Szecsi *et al.*, 2006; Varaud *et al.*, 2011). Relatively little is known about the genetic regulation of petal development in soybeans; however, there are a handful of identified mutants that exhibit altered petal formation. Most of these genes are homologs of known regulators of petal development in Arabidopsis, for example, a MADS-box transcription factor (*GmMADS28*) has been characterized for its role in floral organ specification, the soybean homolog of *LEAFY2* (*GmLFY2*) was found to function in floral organ specification, and a soybean homolog of *UNUSUAL FLORAL ORGANS* (*GmUFO1*) has been implicated in regulating floral organ number and shape (Huang *et al.*, 2014; Wang *et al.*, 2025; Yu *et al.*, 2023).

In many angiosperms, floral organ identity is determined by the ‘ABC’ model: A-class gene expression specifies sepals; overlapping expression of A– and B-class genes specifies petals; B– and C-class gene expression defines stamens; and C-class gene expression alone determines carpels (Bowman *et al.*, 1989, 2012; Coen and Meyerowitz, 1991; Moyroud and Glover, 2017). In addition, E-class *SEPALLATA* (*SEP*) genes are necessary for all four floral whorls to gain proper specification (Colombo *et al.*, 1995; Pelaz *et al.*, 2000).

Though both soybean and *Arabidopsis* have perfect flowers (i.e., they develop all four floral organ whorls), their floral morphology differs. *Arabidopsis* flowers are radially symmetric, and develop floral organs that have more or less equal size and shape. On the other hand, soybean flowers are bilaterally symmetric, which has significant implications for pollinator studies, as bilateral symmetry is generally preferred by bumblebees (Citerne *et al.*, 2010). Unlike *Arabidopsis*, soybean flowers are composed of five sepals, five petals, ten stamens, and a singular carpel with one to four ovules. The petals are further separated into one large standard petal, two wing petals, and two keel petals, each with distinct morphology (Kunst *et al.*, 1989). Despite these differences in floral morphology, most hypotheses regarding soybean floral development remain based on *Arabidopsis* and Antirrhinum (*Antirrhinum majus*) (another bilaterally symmetric floral model) for floral organ specification (Coen and Meyerowitz, 1991; Jung *et al.*, 2012).

One of our goals was to leverage knowledge from *Arabidopsis* to further expand our understanding of the genetic regulation of soybean petal development. In this study, we present a gene expression and open chromatin profiling atlas for reproductive development in soybean. Using a combination of traditional transgenic complementation and new CRISPR activation (CRISPRa) technology (Lowder *et al.*, 2017) we target gene candidates expressed in reproductive tissues to increase soybean petal size (*GmANT* and *GmBPE*), sugar content in nectar (*GmSUC2*), and petal color (*W1*). While our targeted modifications for petal color successfully transformed Williams 82 flowers from white to purple, our attempts to modify petal size proved to be more challenging. In contrast to our predictions based on gene function in *Arabidopsis*, we found that transcriptional activation of *GmANT* and transcriptional repression of *GmBPE* decreased petal sizes. Our results indicate that the genetic architecture for petal development has diverged substantially between *Arabidopsis* and soybean. Our floral organ expression atlas and supporting floral H3K4 trimethylation ChIP-Seq data provide foundational resources for the future targeted modification of additional genetic candidates.

## Results

### Building an enhanced resource for targeted modification of floral traits in soybean

Relative to other plant model systems, soybean lacks genomic resources for identifying gene targets to modify floral organ traits (Ma *et al.*, 2005; Neumann *et al.*, 2022; Schmid *et al.*, 2005). To fill this gap and accelerate reproductive trait modifications in soybean, we constructed a floral organ expression atlas and overlaid these data with open chromatin mapping using H3K4 trimethylation ChIP-Seq. H3K4 trimethylation is used as an epigenetic marker to identify regions of open chromatin around transcriptionally active gene coding sequences (Zhang *et al.*, 2009). Using hand-dissected organs (carpels, stamens, petals, and sepals harvested at early floral organogenesis) we generated a 3’ RNA-Seq expression atlas (Figure 1a). In addition to the four whorls dissected from young buds (≤ 1 mm), we collected entire young floral buds (≤ 1 mm, not showing petals) and mature flowers (with emerged petals) (Figure 1b,c). We also generated young leaf transcriptomes to compare with each floral whorl for the identification of floral organ enriched gene expression patterns.

**Figure 1:**
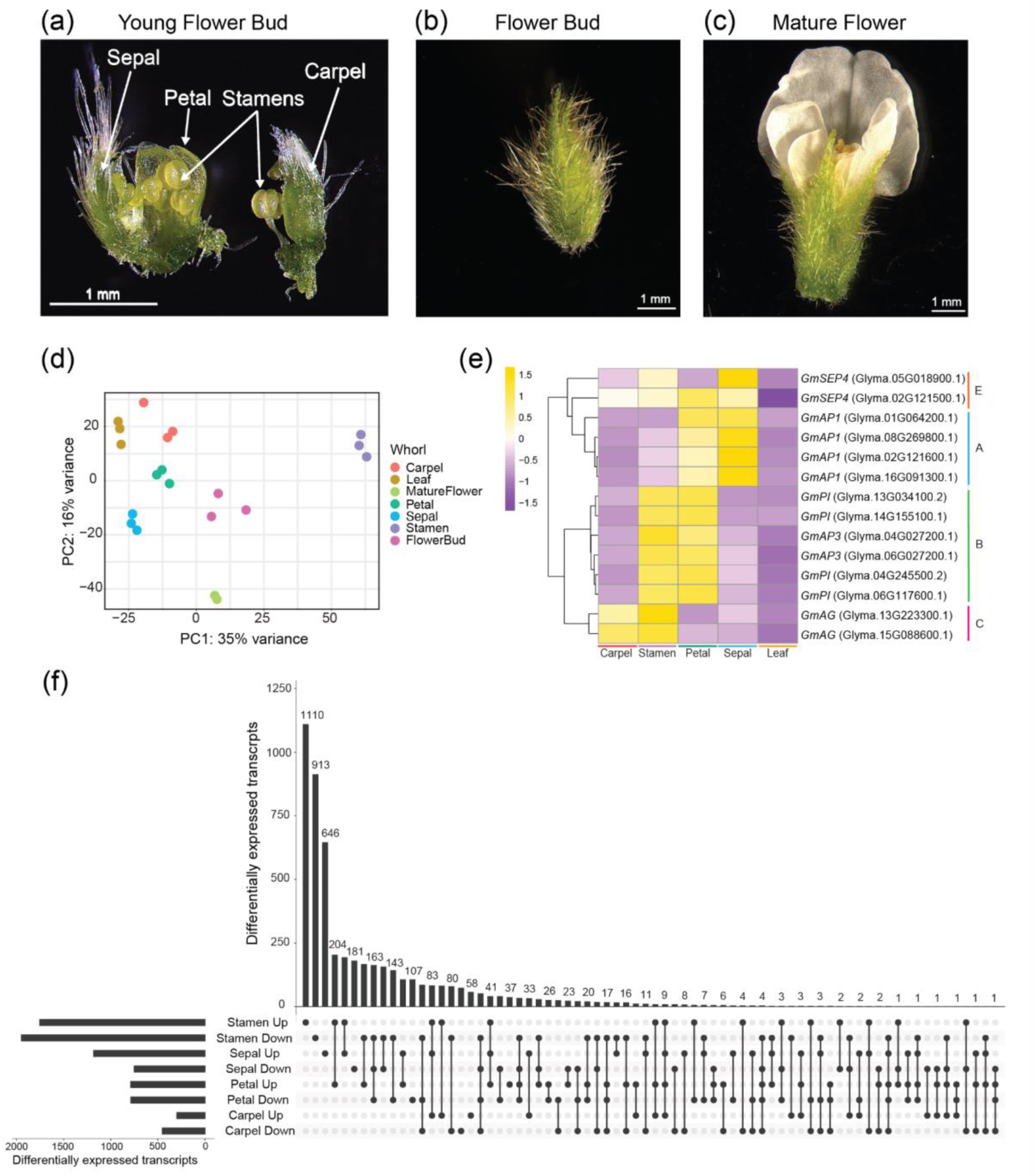
An organ-specific expression atlas for floral development in soybean. The atlas was generated using four hand-dissected floral whorls: sepal, petal, stamen, and carpel (a), floral buds (b), and mature flowers (c). Biological replicates show clean clustering based on PCA analysis (d), and distinct expression profiles. A heatmap (e) of significantly upregulated MADS-box transcription factors in soybean are clustered and scaled by row. The colors correspond with z-score normalization where white indicates the average expression, yellow is an above-average expression and purple is a below-average expression. An upset plot (f) groups the overlap between whorls for differentially upregulated and downregulated transcripts.

Based on gene expression, our biological replicates cluster into clean groups in our PCA (PC1 explains 35% of the variance and PC2 explains 16% of the variance), demonstrating that the organs are expressing distinct transcriptomes (Figure 1d). Using pairwise differential gene expression analysis between each floral organ and leaf tissue, we identified hundreds-to-thousands of significantly differentially expressed genes (DEGs) for each whorl (adjusted p-value ≤ 0.05): 754 DEGs in carpels, 3,704 DEGs in stamens, 1,576 DEGs in petals, and 1,935 DEGs in sepals (Figure 1f). Most of the whorls exhibited partially overlapping expression profiles, with the exception of the stamens, which had over 2,000 distinct DEGs (Figure 1f). As an additional marker of sample quality, we tested the extent to which MADS-box genes in soybean exhibited whorl-specific expression that fits with the well-documented ‘ABC’ model for floral organ specification (Soltis *et al.*, 2007; Theissen *et al.*, 2000; Weigel and Meyerowitz, 1994). Out of 39 identified MADS-box transcription factors, we found that 14 were significantly upregulated in one or more floral whorls compared to leaf samples (Figure 1e). Based on our expression analysis, we can infer that MADS-box genes in soybean exhibit expression patterns that largely align with the ‘ABC’ model (Figure 1e). A-class soybean paralogs of *APETALA1* (*GmAP1*) have an increased expression in petals and sepals; B-class paralogs of *PISTILLATA* (*GmPI*) and *APETALA3* (*GmAP3*) are expressed in stamens and petals; C-class paralog, *AGAMOUS* (*GmAG*), is expressed in carpels and stamens; and E-class paralogs of *SEPALLATA4* (*GmSEP4*) are expressed throughout all the whorls (Figure 1e).

To identify large-scale functional trends in our data, we performed a gene ontology (GO)-term enrichment analysis for significantly upregulated transcripts in each whorl. Our findings are largely congruent with biological expectations. In carpel samples, the top GO terms are associated with ovule/floral organ development and morphogenesis; in stamens, the top terms are associated with micro-gametogenesis; in petals, the top terms are associated with floral organogenesis; and in sepals, there are terms for floral organ formation along GO terms not seen in the other whorls, such as the organic hydroxy compound biosynthetic process and the flavonoid metabolic process (Supporting Figure 1). In addition, we identified overlapping enriched GO terms between neighboring whorls, for instance, stamen formation in the stamen and carpel datasets.

These overlapping terms likely result from B– and C-class gene patterning across neighboring whorls (Supporting Figure 1a,b). We also used pairwise comparisons to identify transcripts that are uniquely upregulated in each organ dataset. Stamens have the greatest number of significantly upregulated unique genes (735), followed by sepals (452), carpels (101), and petals (97) (Supporting Figure 2a-d). When we compared the GO terms for transcripts uniquely upregulated in stamens, we uncovered terms that are specific to anther function, for instance, pollen wall assembly and pollen exine formation (Supporting Figure 2e). Overall, our analyses indicate that this new dataset provides high-quality, organ-specific expression data that aligns with our general expectations for floral patterning from other model systems.

**Figure 2:**
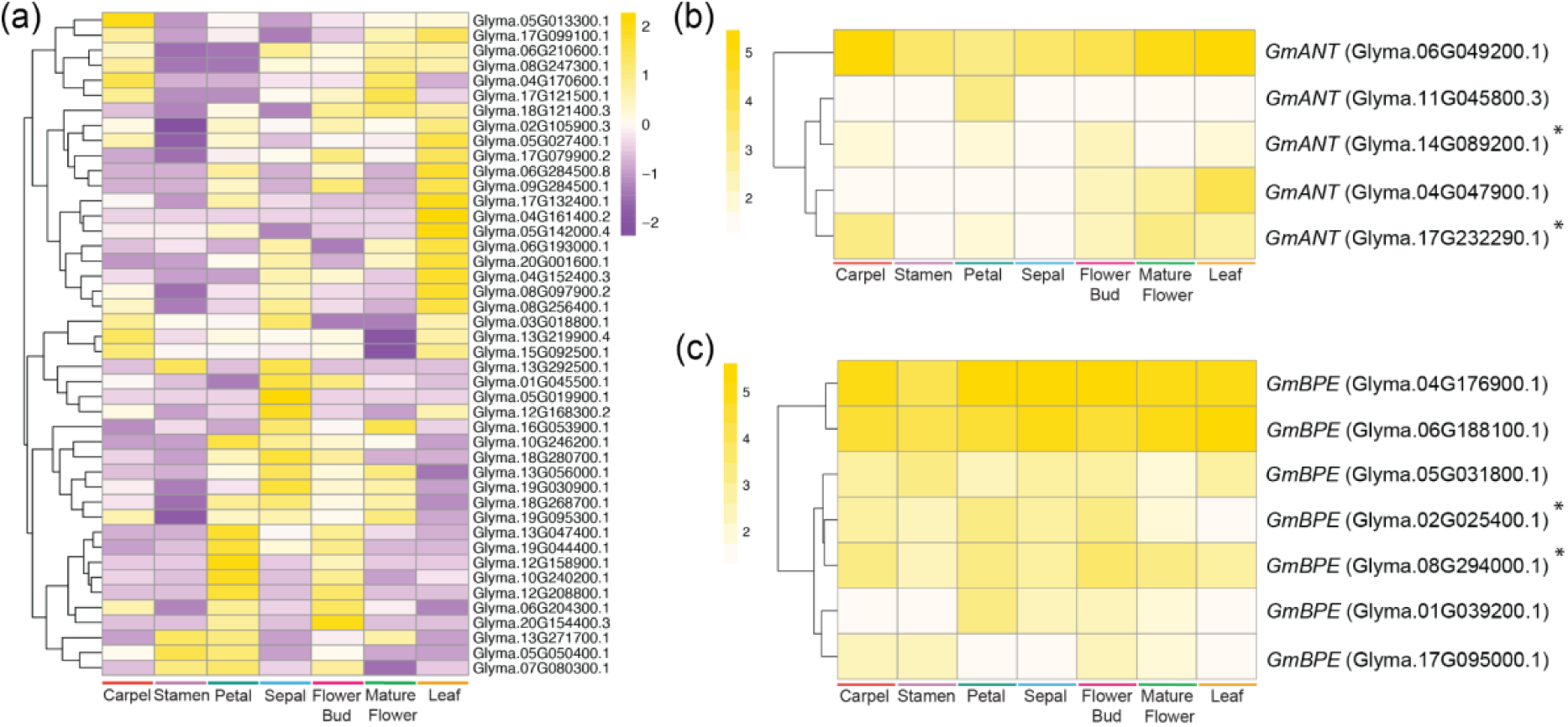
Gene paralogs differentially expressed in various whorls at different expression levels. a) Heatmap of *GmTCP* paralog expression. Heatmap is scaled by row and colors match z-score normalization. Yellow indicates an increase in expression while purple dictates a decrease in gene expression. (b) *ANT* paralogs and (c) *BPE* paralogs in *G.max* have differential normalized counts. Heatmaps are clustered by row and the color yellow indicates higher counts. The paralogs used in this study are denoted with an asterisk.

### H3K4 trimethylation ChIP-Seq of floral buds identifies open chromatin targets for reproductive modification

Unlike traditional CRISPR, CRISPR activation (CRISPRa) and repression (CRISPRi) are used to modulate gene expression without inducing double-strand breaks or permanent edits to a genome. CRISPRa/CRISPRi typically uses a catalytically inactive Cas9 (dCas9) and guide RNAs (gRNAs) to recruit transcriptional activation (e.g., VP64) or repression (e.g., SRDX) machinery to specific gene promoters (Lowder *et al.*, 2017, 2015, 2018; Pan, Sretenovic, *et al.*, 2021; Pan, Wu, *et al.*, 2021; Tang *et al.*, 2017). Alternative strategies using catalytically active Cas9 fused to activation domains with truncated gRNAs have also been developed to achieve transcriptional activation without DNA cleavage (Pan *et al.*, 2022). This technology allows for spatial and temporal regulation of transcriptional activation/repression through the selection of gene promoters that drive Cas9 expression, either deactivated or catalytically active, in a developmentally controlled manner. To aid in the design of CRISPRa gRNAs that target open chromatin regions around our target gene promoters, we performed H3K4 trimethylation ChIP-Seq profiling for two reproductive stages, flower buds and mature flowers, generating genome-wide reproductive floral development transcription factor (TF) binding profiles (Figure 3b-d). Using the high-quality peaks present in all three replicates of full flower buds and mature flowers, we identified peaks for putative open chromatin regions that overlap within 1 kb of 49,792 and 48,629 CDS regions, respectively. We annotated these high quality peaks in the two floral stages of development and determined that 80% of open chromatin regions were within the promoter region and/or within 1kb of transcriptional start sites (TSS) (Supporting Figure 3c,d). When we compared DEGs that were upregulated in our floral organ samples with high quality open chromatin peaks, we found that 70% of whole bud and flower bud DEGs overlap with open chromatin, demonstrating that this H3K4 trimethylation ChIP-Seq dataset generally aligns with transcriptionally active regions of the genome (Supporting Figure 3a,b).

**Figure 3:**
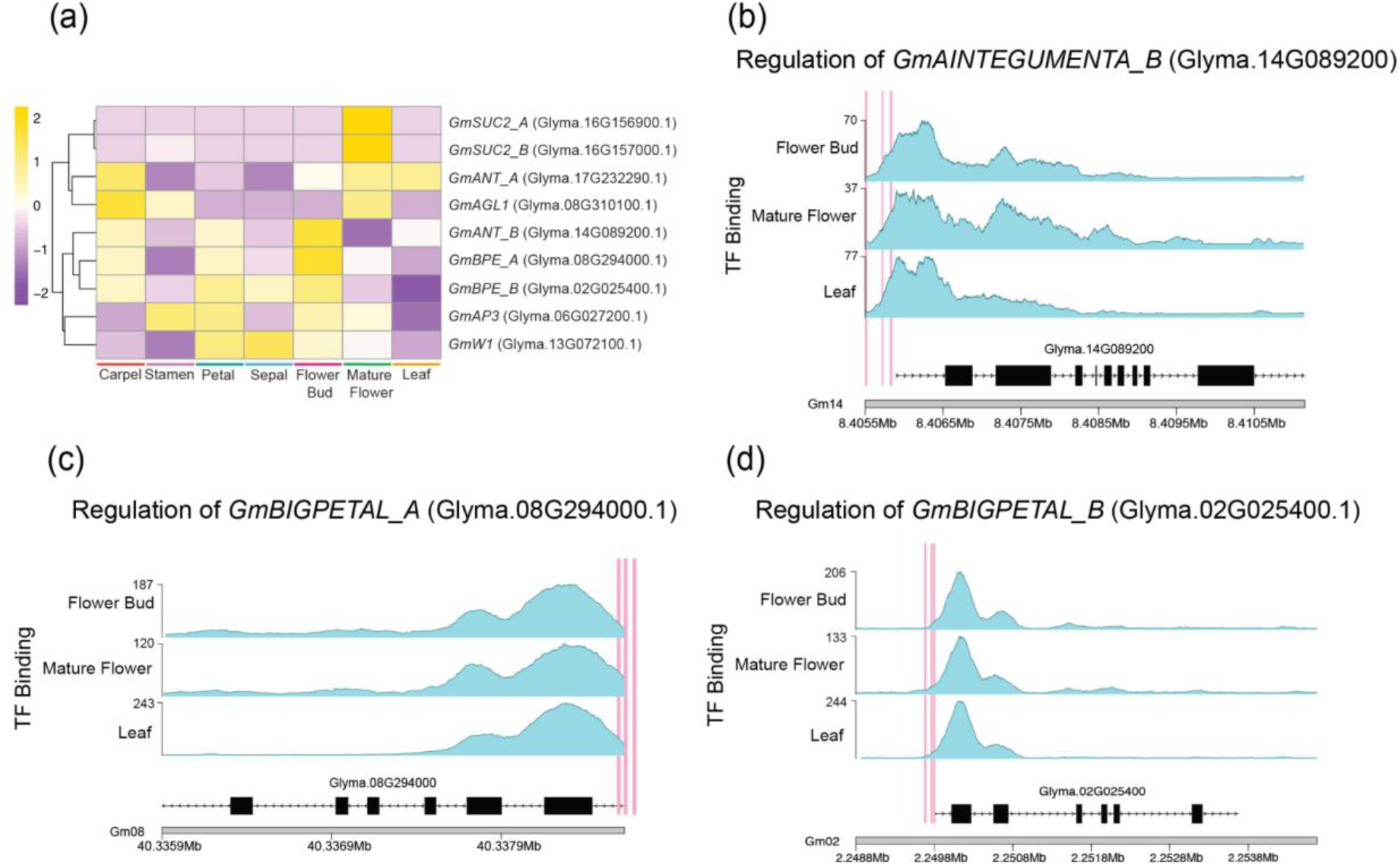
Selection and design of transcriptional gene targeting to enhance pollinator preferences in soybean. (a) Z-score heatmap of the genes targeted in this study are scaled and clustered by row. (b-d) Identified putative TF-binding sites in three tissue types (flower bud, mature flower, and leaf) in target genes. The blue peaks represent regions of open chromatin and the pink line indicates the target sites for engineered gRNAs.

### Genomic and transcriptomic identification of target genes to enhance soybean floral display

With the goal of making soybeans more attractive to bee pollinators, we designed constructs to enhance key pollinator preferences, including increasing petal size, changing petal color, and increasing percent sugars in nectar (Waller 1972; Erickson 1975; Barda et al. 2023). We selected floral target genes based on published studies in *Arabidopsis* and enriched expression of putative soybean orthologs during reproductive development. We identified candidate targets for altering petal size and color by annotating a list of upregulated transcripts in petals, based on differential expression analysis between the petal and leaf transcriptomes. Within this list, we found many MADS-box transcription factors, including A– and B-class genes, *APETALA1* (*AP1*), *PISTILLATA* (*PI*), and *APETALA3* (*AP3*), and E-class genes *SEPALLATA1-3* (*SEP1-3*). We also generated a list of genes specifically upregulated in petals relative to the other three floral whorls and identified 97 petal-specific genes, 86 of which are also present in our petal versus leaf dataset.

Next, we focused on gene families that have demonstrated roles in regulating petal size, including the *TCP* (*TEOSINTE-BRANCHED1/ CYCLOIDEA/ PROLIFERATING CELL NUCLEAR ANTIGEN FACTOR (PCF)*) family, *AINTEGUMENTA* (*ANT*), and *BIG PETAL* (*BPE*). TCP transcription factors regulate organ size by controlling cell proliferation and expansion Of the 55 *TCP* paralogs that we identified in soybean, 44 were expressed in our dataset and 7 were highly upregulated in petals and flower buds (Figure 2a). Given the size and broad expression of the *TCP* family, we decided to focus on other gene families for altering petal size. For *ANT*, we identified six *GmANT* paralogs expressed in our dataset (based on ≥ 10 aligned read per sample) (Figure 2b), and two paralogs that were upregulated in petals, and strongly upregulated in developing floral buds (Glyma.17G232290.1 and Glyma.14G089200.1) (Figure 2b). For *BPE* candidates, seven out of the eight paralogs were expressed in our data. While none of the *GmBPE* genes exhibited petal-specific upregulation, there were several candidates with increased petal and floral bud expression relative to the other samples that we targeted for transcriptional activation (marked with asterisks in Figure 2c). We also searched for sucrose transporters that exhibited high expression during floral maturation that could be targeted for increased percent sugars in nectar. Two candidates from our search, *GmSUC2_A* (Glyma.16G156900) and *GmSUC2_B* (Glyma.16G157000), showed high expression in mature flower samples (Figure 3a). In addition to selecting new candidates for CRISPRa/CRISPRi modification, we also designed a construct to modify petal color from white-to-purple using functional allele complementation with a wild sequence of *W1*, which is known to regulate purple petal color in soybean (Figure 3a).

To restrict transcriptional activation/repression to targeted organs within developing flowers, our CRISPRa/CRISPRi constructs included gRNAs targeting our gene candidates for floral trait modification, as well as deactivated Cas9 (dCas9) that was developmentally regulated by B– and C-class floral organ promoters. *GmAGL1* was selected for enhanced carpel/nectary expression of *GmSUC2_A*/*B* targets, and *GmAP3* was selected for enhanced expression of *GmANT_A/B* and repressed expression of *GmBPE_A/B* in petals (Ma *et al.*, 1991) (Figure 3a). We also searched our targeted gene IDs against our ChIP-Seq list and identified open chromatin regions around the transcriptional start site of *GmBPE_A/B* and *GmANT_B* in both developmental stages and leaves (Figure 3b-d). While open chromatin peaks were identified for *GmSUC2_A/B*, these peaks did not pass our filter for read depth (Supporting Figure 8).

Domesticated soybean has two whole genome duplication events in its recent history, dating back approximately 59 and 10-15 million years ago (Gill *et al.*, 2009; Schmutz *et al.*, 2010; Sherman-Broyles *et al.*, 2017), and thus the genome contains multiple copies of most *Arabidopsis* genes (referred to as paralogs throughout this paper). Therefore, we generated two constructs for each soybean paralog for our activation lines, and multiplexed gRNAs on one construct targeting both paralogs for our transcriptional repression lines (Supporting Figure 4a; 7b). Putative soybean orthologs for *BIGPETAL* (*GmBPE*) and *AINTEGUMENTA* (*GmANT*) were selected to increase petal size, based on previous research knockouts that reported a significant change in petal size in *Arabidopsis* and the localized expression of these genes in soybean floral buds (Mizukami and Fischer 2000; Randall et al. 2015; Szecsi et al. 2006; Varaud et al. 2011).

**Figure 4:**
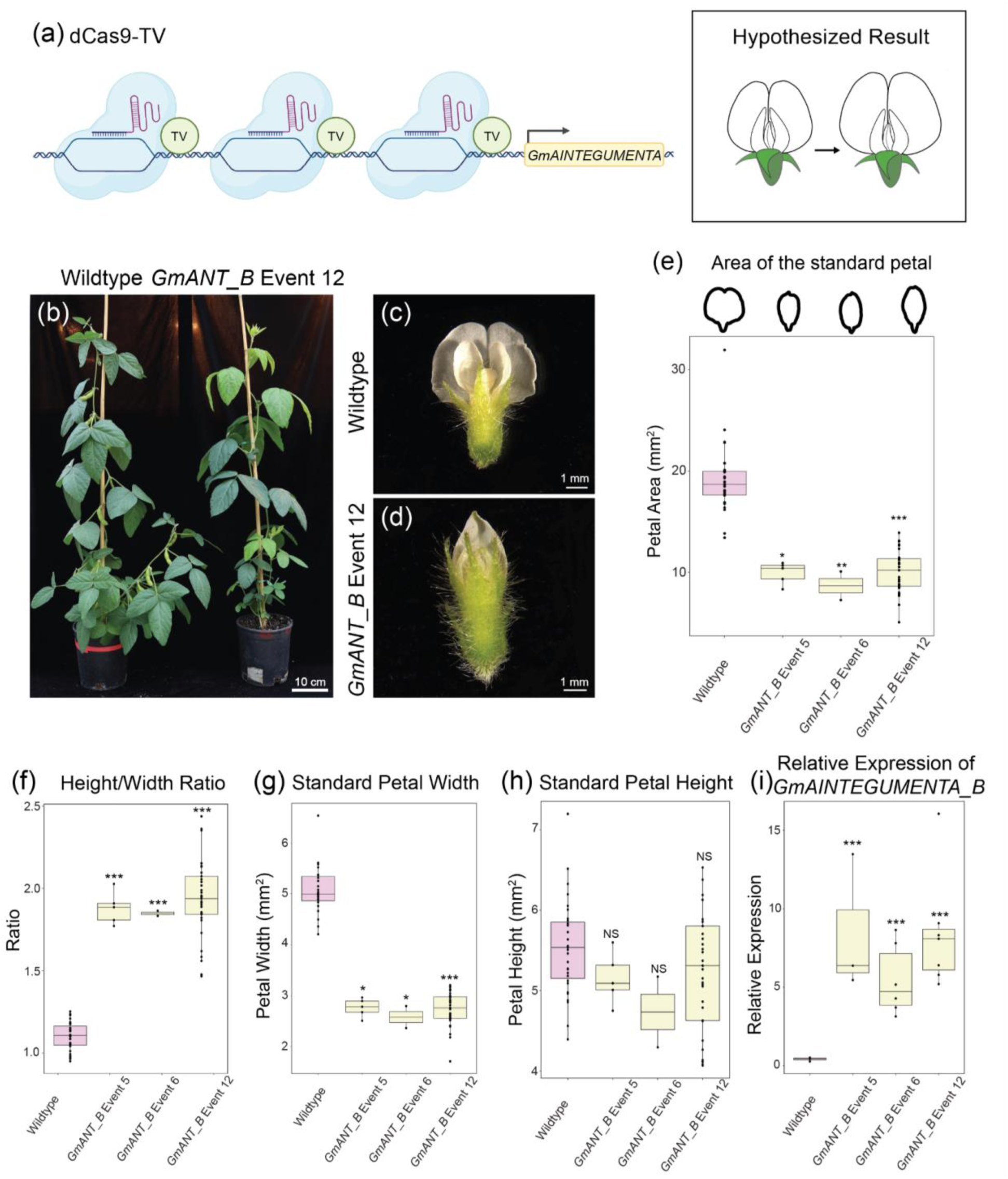
Increased expression of *GmANT_B* decreases petal width. A diagram of our hypothesized results based on *ANT* function in Arabidopsis: increasing *GmANT* expression using CRISPRa would lead to increased petal size (a). Side-by-side comparison of vegetative plant phenotypes between wildtype and GmANT Event 12 plant #12 (b). A comparison of wildtype (c) versus GmANT Event 12 (d) floral morphologies. Boxplot of the area of the standard petal (outline featured above plot) between wildtype, GmANT Event 5, GmANT Event 6, and GmANT Event 12 (e). Boxplots of petal height/width ratio (f), petal width (g), and height of the standard petal (h). To compare petal size, samples were fitted to a linear mixed effect model using the lmerTest package (Kuznetsova *et al.*, 2017) and the p-value was calculated using a Dunnett’s test using the emmeans package (Length, 2025) in RStudio (Posit team, 2023). qRT-PCR data for the relative expression of *GmANT_B* events 5, 6, and 12 versus wildtype floral buds (i). Since the variance of the qRT-PCR data was significant (according to a Bartlett test conducted using the R stats package (R Core Team, 2023) in RStudio) the log of relative expression was fitted to a linear model using ANOVA from the R stats package and the p-value was calculated using a Dunnett’s Test. *** = p < 0.001; ** = p < 0.001; * = p < 0.01.

### CRISPR activation of soybean floral targets leads to unexpected effects on petal morphology and size

*Arabidopsis aintegumenta* (*ant*) loss-of-function mutants develop flowers with smaller petals, while *AINTEGUMENTA* overexpression increases cell proliferation and produces larger floral organs (Mizukami and Fischer, 2000; Randall *et al.*, 2015). Based on this, we hypothesized that activating *GmANT* expression in soybean petals would increase petal size. To test this, we used a CRISPRa system dCas9-TV that uses a dCas9 fused to transcriptional activation machinery, which contains six TAL effector repeats and two VP64 domains (TV) known to promote robust transcriptional activation in plants (Figure 4a) (Li *et al.*, 2018; Xiong *et al.*, 2021).

We constructed two vectors to activate soybean *ANT* paralogs, *GmANT_A* and *GmANT_B*. In each, we programmed three gRNAs to recruit dCas9-TV to regions within 200 bp upstream of the transcriptional start sites (TSS) of *GmANT_A* and *GmANT_B* (Figure 3b-d; Figure 4a; Supporting Figure 4a). To repress *GmBPE* paralogs, we expressed dCas9-SRDX under the control of B-Class *proGmAP3* promoter, thereby restricting repression to petals and stamens. We also generated a reporter line (*proGmAP3:dCas9-TV::GUS-tdTomato*) to validate *GmAP3* promoter activity in floral petals and stamens (Supporting Figure 6).

We analyzed more than 10 independent transformation events for each *GmANT_A* and *GmANT_B* CRISPRa (dCas9-TV) construct, focusing on changes in petal morphology. While we did not observe any obvious phenotypes in the *GmANT_A* transcriptional activation lines, we did observe significantly decreased petal size in select *GmANT_B* events that had significantly increased expression of *GmANT_B* (Figure 4c,d). In addition, we observed altered vegetative phenotypes including stunted growth, curled leaves, and dark green venation across multiple plants in these same events (*GmANT_B* events 5 and 12, two thirds of the plants in event 6) (Supporting Figure 4b-d). Surprisingly, instead of increasing the petal area as hypothesized, transcriptional activation of *GmANT_B* reduced the width of the standard petal by approximately 50%, resulting in a narrow petal (Figure 4c-h). To determine whether petal allometry had been altered, we measured the length and width of the standard petal and found that transcriptional activation of *GmANT_B* led to a significant shift in petal shape, with an increased length-to-width ratio (a mean of 1.1 in wildtype vs. 1.8 in transgenic lines) (Figure 4f). Specifically, we found that reduced petal size in our transgenic lines was attributable to a significant decrease in petal width while length remained constant (Figure 4g,h). To confirm that altered petal size correlates with increased *GmANT_B* expression, we performed qRT-PCR and found that *GmANT_B* expression in young flower buds was significantly higher in the transcriptional activation lines with altered petal morphology compared to wildtype controls (Figure 4i). Moreover, we demonstrated that this phenotype persisted across multiple greenhouse trials, indicating that transcriptional activation is stable in these lines (Supporting Figure 4e-g).

To test whether altered cell size contributes to reduced petal width in *GmANT_B* lines, we stained the standard petal of our transcriptional activation and wildtype control lines with calcofluor white and used confocal microscopy to image the epidermal cells on the adaxial side of the petal. We used MorphoGraphX (Barbier de Reuille *et al.*, 2015) to measure the area of ∼200 cells per replicate (Supporting Figure 5a,b). We observed a relatively narrow range of average cell areas in our wildtype replicates (349-361μm^2^/cell). While the *GmANT_B* transcriptional activation lines exhibited more variability in cell size, with average areas ranging from 275-371 μm^2^/cell, we did not observe a significant difference in average cell size between *GmANT_B* petals and wildtype (Supporting Figure 5c).

**Figure 5:**
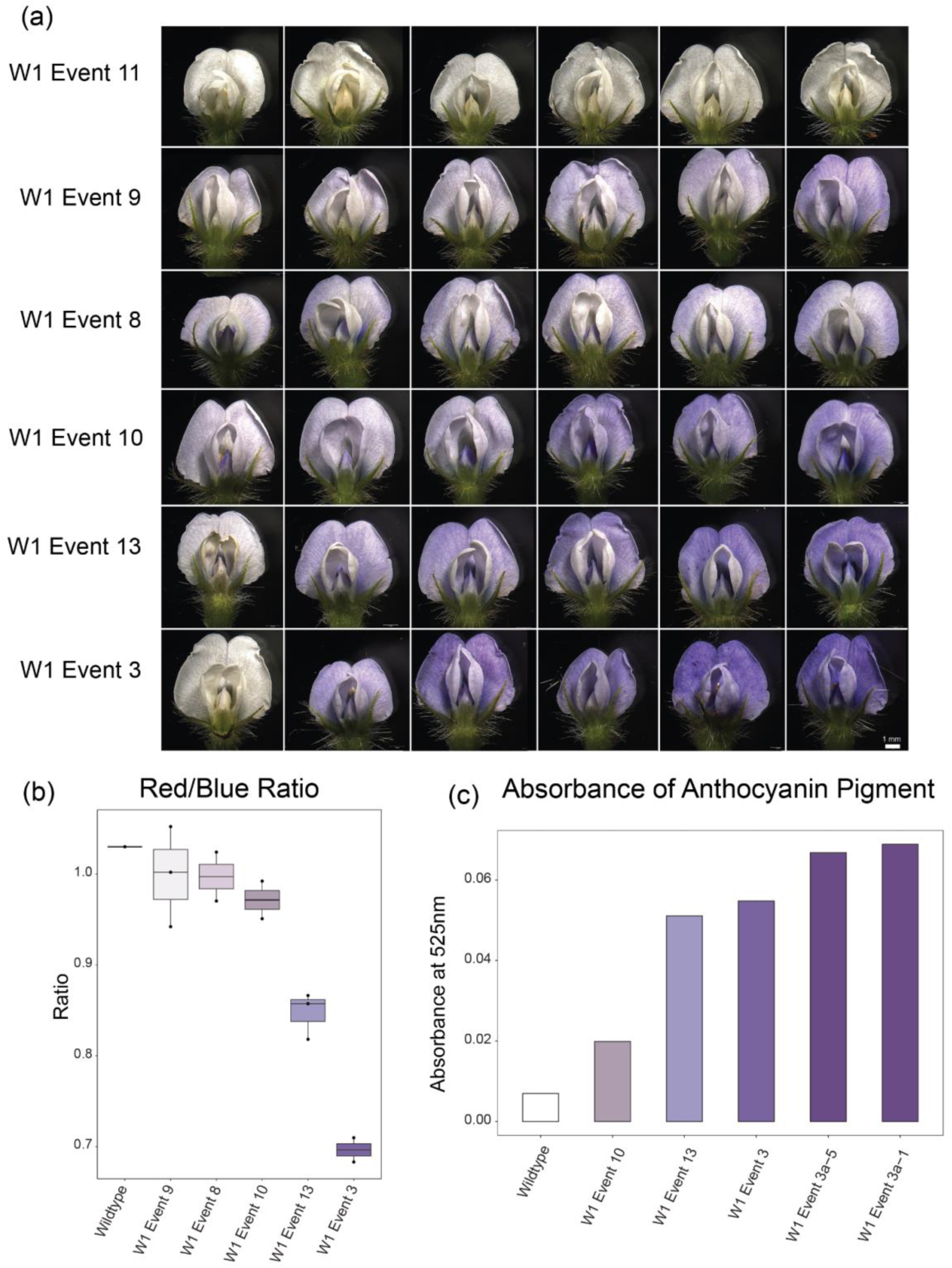
Expression of *W1* in Williams 82 turns on anthocyanin production in soybean petals. Flowers from six independent T1 *W1* events displaying a range of purple petal phenotypes (a). Each flower was harvested from an independent transgenic plant within a transgenic event carrying a functional *W1* transgene. Red/Blue color ratio negatively correlates with the degree of purple petal pigmentation, providing a simple quantification for petal phenotypes (b). Spectrophotometer quantification for anthocyanins (absorbance measured at 525 nm) demonstrates that Event 3 has the highest level of anthocyanin accumulation, which is further boosted in the T2 generation (W1 Event 3a-5 and Event 3a-1) (c).

For our second putative regulator of petal size, we evaluated 10 independent events for our CRISPRi construct targeting *GmBPE_A/B*. Since *BPE* limits petal cell size in *Arabidopsis*, we hypothesized that fusing transcriptional repressor, *SRDX*, to dCas9 would decrease the expression of *GmBPE*, resulting in increased cell proliferation and an overall increase in petal size (Lowder *et al.*, 2017; Szecsi *et al.*, 2006). To guide the dCas9-SRDX protein to the promoter region of *GmBPE*, we designed three gRNAs for each of the two paralogs within 200 bp upstream of the transcriptional start site (Figure 3c,d; Supporting Figure 7a). In order to simultaneously repress both paralogs, we included all six gRNAs in a single construct and expressed the dCas9-SRDX protein fusion under the B-Class gene promoter, *proGmAP3,* restricting transcriptional repression of *GmBPE_A* and *GmBPE_B* to targeted reproductive organs (Supporting Figure 7b). In addition, we created a promoter-reporter line (*proGmAP3:dCAS9-TV:GUS-tdTomato*) to confirm the localization of *GmAP3* expression in floral petals and stamens (Supporting Figure 6). We screened 14 independent transgenic events for altered petal size (based on the area of the standard petal) and a significant reduction in *GmBPE_A* and *GmBPE_B* transcript abundance relative to WT controls. Similar to our results for *GmANT_B* CRISPR targeting and in contrast to our initial expectations, we identified three events that exhibited reduced petal size (GmBPE Event 6, GmBPE Event 9, and GmBPE Event 14) (Supporting Figure 7c). While qRT-PCR data of young flower buds showed a trend towards reduced expression of *GmBPE_A* and *GmBPE_B* paralogs between wildtype and GmBPE events 6, 9, and 14, this difference was not statistically significant, indicating that this construct was less effective than the *GmANT_B* construct (Supporting Figure 7d,e).

To upregulate *GmSUC2* expression in soybean nectaries for increased percent sugar in nectar, we expressed dCas9-TV under the C-class gene promoter, *AGAMOUS-LIKE1* (*GmAGL1*), and designed three gRNAs for each of the two *GmSUC2* paralogs within 200 bp upstream of the *GmSUC2* transcriptional start site (Supporting Figure 8a). Unfortunately, these transgenic lines did not significantly alter the expression of *GmSUC2* (Supporting Figure 8b). In addition, we found that our wild type and negative transgenic control lines exhibited high levels of variability for sugar concentrations in extracted nectar based on extraction time and greenhouse conditions (Supporting Table 1). These data indicate that while our CRISPRa approach was effective at modifying floral morphology, the current design was insufficient to alter *GmSUC2* expression.

### Functional *W1* is sufficient to transform soybean flowers from white to purple

In addition to larger floral displays and sweeter nectar, bee pollinators generally prefer petals that reflect light in the UV/blue region of the color spectrum (Giurfa *et al.*, 1995; Leleji, 1973). We used a naturally occurring, functional allele for *W1*, a flavonoid 3’, 5’-hydroxylase involved in anthocyanin production, to test whether we can transform Williams 82 from a white to purple petalled variety that would be more attractive to bee pollinators. By expressing the coding sequence of *W1* derived from a purple flowered wild relative of *Glycine max* (*Glycine soja*) behind the native *w1* promoter from Williams 82 (Supporting Figure 9a), we were able to transform Williams 82 into a purple flowered variety (Figure 5). Notably, the degree of pigmentation varied by transgenic event in the T1 generation (Figure 5a). We used two methods to quantify pigmentation across transgenic events: first, we measured the red/blue ratio of scanned petals (darker purple petals correspond to lower red/blue ratios, Figure 5b), and second, we measured the absorbance of extracted anthocyanins at 525 nm (Figure 5c). Transgenic event #3 stood out as exceptionally high in anthocyanins relative to the other transgenic lines. Notably, we found that anthocyanin concentrations could be further boosted in the T2 generation, which is likely due to a higher percent of homozygous vs hemizygous transgenic individuals, with *W1* thus showing additive effects or incomplete rather than complete dominance in *Glycine max* (*W1* Event 3a-5 and Event 3a-1) (Figure 5c).

To test whether pollinators prefer purple over white petals in soybean, we designed a choice test in which a plant from the *W1* purple flowered Event 3 was placed next to white flowered Williams 82. The plants were sealed in a tent with mosquito netting and bumblebees (*Bombus impatiens*) were released one at a time into the tent (Supporting Figure 9b). Because the bumblebees continued to forage on whichever plant they visited first, we scored color preference based on the first flower that each bumblebee visited rather than total visitations (Pyke, 1978). We originally performed these trials with honeybees; however, after a failed trial in which the honeybees tried to escape the cages rather than forage on the plants, we switched to bumblebees, which are generally more amenable to greenhouse trials (Supporting Figure 9c). We performed two replicate trials for bumblebee color preference and found that the pollinators trended toward visiting the purple flowered transgenic line over white flowers (59-67% preference for purple over white flowers) (Supporting Figure 9d,e), however, this difference was statistically insignificant based on a one-tailed binomial test (trial 1: p = 0.221; trial 2: p = 0.061) with a sample size of 27 trials in each experiment (Supporting Figure 9d,e).

## Discussion

Soybean has yet to benefit from a high-throughput approach to hybrid breeding. In our previous work, we developed a male-sterility/male-rescue system that could be used to enforce outcrossing in soybean (Szeluga *et al.*, 2023). Our goal for this study was to recruit bee pollinators through enhanced floral traits; these pollinators could then be used to perform hybrid crosses at scale. Using a combination of traditional transgenic complementation and CRISPR activation (CRISPRa) technology (Lowder *et al.*, 2017) we targeted gene candidates expressed in reproductive tissues to increase soybean petal size (*GmANT* and *GmBPE*), sugar content in nectar (*GmSUC2*), and petal color (*W1*).

Using a targeted transcriptional activation/repression approach, we successfully increased the expression of *GmANT_B* mRNA in young flower buds using dCas9-TV, however, this had the opposite effect on the petal phenotype than we had originally hypothesized based on *ANT* function in *Arabidopsis*. Previous studies have shown that knocking out *ant* function produces flowers with a reduction in petal size and petal cell number and overexpressing *ANT* leads to larger organs in *Arabidopsis* (Randall *et al.*, 2015). However, we found that enhancing *GmANT_B* expression in soybean led to flowers with reduced petal sizes. In *Arabidopsis*, *ANT* is directly linked to 200 downstream target genes based on ChIP-Seq experiments, making it a core transcriptional regulator during reproductive development (Krizek *et al.*, 2020, 2016). Evolutionary divergence between soybean and *Arabidopsis* has likely led to rewiring of these downstream transcriptional targets, complicating the impact that *GmANT_B* transcriptional modification has on floral organ development. For *GmBPE_A/B,* we saw a reduction in *GmBPE_A/B* expression compared to wildtype, however, this reduction was not statistically significant. Similar to what we observed for *GmANT_B*, *GmBPE_A/B* events that expressed significant changes to petal size all resulted in smaller petals, which again, is the opposite of what we hypothesized based on *Arabidopsis BPE* function (Supporting Figure 7c). An improved construct design, wherein the gRNAs align more closely with the open chromatin peaks in our ChIP-Seq data, should enhance transcriptional repression at the *GmBPE* loci, resulting in a more significant change in petal morphology. Based on our analysis of CRISPRa of *GmANT* and CRISPRi of *GmBPE_A/B*, we think an informative future experiment would be to reverse these transcriptional modifications (i.e., perform transcriptional repression of *GmANT_B* and transcriptional activation of *GmBPE_A/B*). This experiment would resolve if petal size can be directly modified through transcriptional control of these target genes, or whether there are more complex regulatory interactions that need to be considered to successfully engineer this trait.

We successfully introduced anthocyanin production in the petals of the white-petal cultivar, Williams 82, and saw a variation of purple hue within and across transformation events. The increase of anthocyanin absorption of the T2 generation of W1 Event 3 compared to the T1 generation suggests that the anthocyanin concentration and degree of purple hue are due to the dosage of *GmW1* expression in the petals. When we compared the attractiveness to bumblebees of our purple lines to Williams 82, we saw a preference for purple petals in two separate trials. An additional experiment with a larger sample size may yield significant results; however, conducting these trials is a resource-intensive endeavor, as both plants and pollinators need to be in optimal health at the time of the trial and trials are time-intensive to monitor. Even though some soybean varieties already produce anthocyanin pigmentation, it is helpful to be able to engineer this trait in white flowered varieties without relying on breeding. Not every cross results in heterosis; the efficiency of hybrid breeding is highly dependent on the parental varieties being crossed and these parents may naturally have white petals. In addition, existing cultivars with naturally purple color may be genetically linked to an undesirable phenotype to breeders while a white-colored cultivar is superior (Naves *et al.*, 2022). Our demonstration that soybean petal color can be transgenically altered (and thus also potentially gene edited) provides a useful tool for controlling this trait.

In addition to demonstrating that floral traits can be altered through targeted modifications, we have created a floral whorl atlas that will serve as a valuable resource for investigating reproductive development in soybean. The clear separation of our samples by group in the PCA plot and the arrangement of GO term functions by whorl support the quality of our RNA-Seq data. We paired the ChIP-Seq data with the RNA-Seq data to identify regions of open chromatin accessible for transcription and gene expression. Expression of key regulators, the MADS-box domain transcription factor family, is functionally conserved in soybean and these maintained localized expression within the standard floral ‘ABC’ model.

A single gene in *Arabidopsis* has multiple orthologs in soybean due to the multiple rounds of whole genome duplication (Schmutz *et al.*, 2010; Shoemaker *et al.*, 2006). *Arabidopsis* has a total of five chromosomes and approximately 27,000 coding genes (*Arabidopsis* Genome Initiative 2000), while soybean has 20 chromosomes derived from at least two genome duplication events resulting in approximately 64,000 protein-coding genes (Severin et al. 2010; Walling et al. 2006). As a result of evolution, these duplication events have led to divergent gene functions between *Arabidopsis* and soybean, which are evolutionarily separated by ∼90 million years (Grant *et al.*, 2000). When fully characterized, having differentially expressed paralogs can be beneficial for designing dosage-dependent constructs (Szeluga *et al.*, 2023). Without functional data in soybean it is challenging to accurately predict the phenotypic outcome of targeted gene modifications in this crop species based on *Arabidopsis* gene function alone. As we have learned from our experiments to enlarge petals and increase the percent sugars in nectar, relying on orthologous expression data in *Arabidopsis* is insufficient to genetically engineer soybeans.

Using our new floral atlas, we were able to identify additional gene families that could be used to alter petal shape and size, including the *TCP* gene family. *TCP* transcription factors are widely involved in plant growth and development. In *Arabidopsis*, *TCP* proteins have been associated with circadian rhythm, stress response, plant morphology, floral asymmetry, hormone biosynthesis, and cell growth (Camoirano *et al.*, 2023; Cubas *et al.*, 1999; Li, 2015; Li *et al.*, 2021). *TCP1*, specifically, is involved in the brassinoid biosynthesis pathway and plant morphology, acting as a negative regulator for cell division and growth (An *et al.*, 2011; Guo *et al.*, 2010; Li, 2015; Li *et al.*, 2005). Similar to MADS-box transcription factors, the *TCP* protein family is functionally conserved across angiosperms making the translation between species a viable option (Feng *et al.*, 2018; Martín-Trillo and Cubas, 2010). Overexpression of *TCP1* and other *TCP* proteins have led to smaller flowers in *Arabidopsis* (Busch and Zachgo, 2007; Huang and Irish, 2015). Future studies targeting *TCPs* could be informative for modifying petal size and understanding organ size regulation in soybean.

In conclusion, our study demonstrates the challenges and promise of using targeted CRISPRa/CRISPRi to modify floral traits in soybean. While the outcomes of manipulating *GmANT_B* and *GmBPE_A/B* expression deviated from our original hypotheses that were based on *Arabidopsis*, they underscore the importance of performing functional validation to translate basic knowledge into crop systems. Our successful engineering of anthocyanin pigmentation in white-flowered cultivars provides a practical tool for altering petal color in soybean, and our floral whorl expression atlas along with open chromatin profiling, offer foundational resources for future trait modification. Together, these efforts highlight both the potential and complexity of translational genomics in soybean and emphasize the need for species-specific approaches when utilizing model plant data to guide crop improvement.

## Materials and Methods

### Construct design

Transcriptional activation lines in soybean were constructed for *ANT* and *SUC2* paralogs and a single transcriptional repression line for two *BPE* paralogs (Supporting Figures 4,7,8). To promote transcriptional activation, a deactivated Cas9 (dCas9) was fused to transcriptional activator, TV, which is composed of 6 x TAL activation domains and 2 x VP64. The single transcriptional repression line was constructed by fusing dCas9 to an SRDX domain. Target gene expression data for the genes in open-flower and closed-flower were obtained from databases such as Soybase (Brown *et al.*, 2021; Severin *et al.*, 2010), Soyatlas (Almeida-Silva *et al.*, 2023), and Phytozome (Goodstein *et al.*, 2012). To avoid off-target effects from the dCas9 technology, the expression system was driven by a promoter specific to petals (*GmAP3)* for *ANT* and *BPE* and a promoter that is specific to carpels and stamen (*GmAGL1*) for *GmSUC2*. Three gRNAs per gene were designed within the first 200 bp upstream of exon 1. To make the flowers purple, the native white-flowered *Gmw1* promoter was used to drive the purple-flowered *W1* allele. Controller lines were created for promoters *GmAP3* and *GmAGL1* with the addition of GUS-td_tomato to visualize the expression of these promoters in the soybean flower (Supporting Figure 6). The Wisconsin Crop Innovation Center (WCIC) was contracted to build these construct designs and transform them into *Glycine max* cv. Williams 82.

### Plant transformation

Transgenic soybean lines were generated at the WCIC using a proprietary protocol that involves *Agrobacterium tumefaciens*-mediated transformation of Williams 82 cultured shoot apical meristems. Positive transformants were selected based on spectinomycin resistance and confirmed with transgene-specific primers.

### Measuring petal size for GmANT_B and GmBPE events

In 2023, plants were grown at the Donald Danforth Plant Science Center (Olivette, MO) and imaged using a Nikon D7200 camera with AF-S DX NIKKOR 18-140 mm f/3.5-5.6G ED VR lens. Soybean flowers were dissected and flattened during imaging under a clear plastic sheet with a weight on top. Using FIJI (Schindelin *et al.*, 2012), the mid-rib of the back petal was measured to represent the length of the flower while the width was measured as the distance of the widest petal edges, perpendicular to the length. Samples were fitted to a linear mixed effects model using lmer (Kuznetsova *et al.*, 2017) and the p-value was calculated using a Dunnett’s test (Length, 2025).

In 2024, plants expressing the transgene to repress *BPE* were grown in the greenhouse at the Donald Danforth Plant Science Center and imaged using the Zeiss Axio Zoom dissecting microscope with a PlanNeoFluar Z 1x/0.25 FWD 56 mm lens and camera.

Images were analyzed using PlantCV (Gehan *et al.*, 2017). Samples were fitted to a linear mixed effects model using lmer and the p-value was calculated using a Dunnett’s test.

Plants expressing the transgene to increase *GmANT* expression were grown in the greenhouse at Cornell University and were used to calculate petal size. The back petal was dissected, flattened using glass slides, and imaged using a Leica M205 FCA dissecting microscope. Samples were fitted to a linear mixed effects model using lmer and the p-value was calculated using a Dunnett’s test.

### Measuring cell size for GmANT_B *events*

Plants were grown in greenhouse conditions at Cornell University. After petals were imaged to calculate the petal size, they were fixed in FAA overnight, followed by a dehydration series to 70% EtOH. The petal was placed facing up on a glass slide and two drops of 0.1% Calcofluor White and one drop of 10% KOH were placed on the petal to stain the cell walls. The stained petal was pushed flat with a coverslip and immediately excited at 405 nm on a Zeiss u880 confocal microscope at 20x magnification.

A series of 5 tiles 425 micron square were imaged horizontally across the petal, centered at the middle of the petal designated by a series of longitudinal xylem and continuing through the widest width of the petal. Cell size was measured using MorpoGraphX (Barbier de Reuille et al. 2015) software for ∼200 cells per plant and averaged together. Cells partially cut off by the border were not included in the count.

### qRT-PCR quantification of gene expression in CRISPRa events

Young developing flower buds (<1 mm, 15 buds per plant) were collected in the greenhouse, flash frozen in liquid nitrogen, and ground up in a TissueLyser II Bead Mill Sample Disruption Preparation Machine (Qiagen, Hilden, Germany) for 1 minute and 30 seconds, flipping the samples halfway, and extracted using TRIzol (Ambion, Austin, TX, U.S.A). RNA samples were checked for quality using 260/280 and 260/230 ratios on the DeNovix Spectrophotometer and quantified using Qubit^Ⓡ^ RNA Broad Range (BR) Assay (Invitrogen, Waltham, MA, U.S.A) on a DeNovix Spectrophotometer.

For “BIGPETAL*”* lines, 500 ng of RNA was converted to cDNA using Maxima™ H Minus cDNA Synthesis Mastermix with dsDNase (Thermo Fisher Scientific, Waltham, MA, U.S.A) and quantified on a QuantStudio™ 5 Real-Time PCR System, 384-well machine using GoTaq(R) qPCR Master Mix (Promega, Madison, WI, U.S.A). Samples were fitted to a linear model using ANOVA and the p-value was calculated using a Dunnett’s test.

For “AINTEGUMENTA” lines, 500 ng of RNA was converted to cDNA using QuantiTech^Ⓡ^ Reverse Transcriptase (Qiagen, Hilden, Germany). The cDNA was diluted as used with PowerUP SYBR Green Master Mix (Applied Bioscience) to quantify the expression of *ANT_B* using a Roche 480 qPCR machine (Roche Holding AG, Basel, Switzerland). Housekeeping genes UKN1 and FLD were used for delta Ct analysis. For statistical analysis, samples were concluded to have a significant variance using a Bartlett Test and were log-transformed to make the variation non-significant. Samples were fitted to a linear model using ANOVA and the p-value was calculated using a Dunnett’s test.

### GUS staining of *GmAP3* reporter line

Various bud and flower stages were taken from wildtype and *proGmAP3:dCas9-TV::GUS-tdTomato* plants grown in the greenhouse. Tissues were harvested in 90% ice-cold acetone and kept on ice until the remainder of the samples were collected. The samples were vacuum infiltrated at room temperature for 10 minutes, followed by an additional 10 minutes of rest at room temperature. The acetone was removed, and samples were washed with the staining buffer (50 mM Sodium Phosphate Buffer pH 7.0, 10 mM EDTA pH 8.0, 1 mM Potassium Ferricyanide, 1 mM Potassium Ferrocyanide, 0.2% Triton X-100). This staining buffer was removed and replaced with a fresh staining buffer that contained 1 mM of X-Gluc. Samples were incubated at 37°C for two hours. After incubation, the samples underwent a dehydration series and stored in 70% EtOH at 4°C.

### Measuring sucrose concentration in *GmSUC2* flowers

Flowers from plants grown in the greenhouse at the Donald Danforth Plant Science Center were cut at the pedicel at 8:30 AM and used for nectar extraction. The flower was placed face down on a dissecting microscope and a razor blade was used to cut a small opening in the bottom of the flower where the nectary is. Pressure was gently applied to release nectar from the flower. The liquid was taken up via capillary action using a Drummond 0.5 μl MicroCapillary tube (Drummond Company, Birmingham, AL, U.S.A). The nectar was deposited into a PCR tube for collection. Multiple flowers (20 > x > 10) were used to collect enough nectar for a reading. 1.2 μl of nectar was deposited into a new tube with 1.2 μl of distilled water. The sample was mixed and vortexed, and 2.2 μl was added to the surface of a low-volume Eclipse Refractometer (Bellingham + Stanley, Code 45-81) to measure percent sugars in the nectar. The percentage was doubled to adjust for the dilution. Samples were taken from wildtype and control (WP_1283) plants.

### Imaging *W1* petal color

Mature flowers were imaged using a Leica M205 FCA dissecting microscope. Red and blue values were extracted from the images to calculate the red/blue ratio using FIJI (Schindelin *et al.*, 2012).

### Extraction of Anthocyanins from *W1* lines

To quantify anthocyanins, soybean flowers were flash frozen in liquid nitrogen, ground with a chilled mortar and pestle, and suspended in an extraction buffer (1M HCL in 80% methanol to a concentration of 1%). The solution was shaken overnight at 4°C and spun down at 10,000 rpm for five minutes. The supernatant was aliquoted into a UV-transparent cuvette and the OD was measured at 525 nm using a spectrophotometer.

### 3’ RNA sequencing

Floral buds, mature flowers, and leaves were removed from the plant and snap frozen in liquid nitrogen. In addition, young floral buds (< 1 mm) from *Glycine max* cv. Williams 82 were collected and placed in a 1.5 mL microcentrifuge tube filled with 100% acetone in a –20°C freezer block to fix the tissue before dissection. The buds were dissected on a – 20°C freezer block topped with a cleaned, Rnase-zapped™ surface (Thermo Fisher Scientific, Waltham, MA, U.S.A). Individual whorls were dissected with Dumont #5 forceps under a Leica M205 FCA fluorescent stereo microscope and placed into 1.5 mL microcentrifuge tubes containing acetone in another –20°C freezer block. Forceps and the dissecting surface were disinfected and sprayed with RNase-Zap™ between samples. Tubes were spun down, and the acetone was removed with a micropipette. The tubes were snap-frozen in liquid nitrogen, and all tissues (individual whorls, whole buds, mature flowers, and leaves) were ground up with a cold micropestle. RNA was extracted using the RNeasy® Plant Mini Kit (Qiagen®, Hilden, Germany). RNA was cleaned by adding 0.1 volumes of 3M Sodium Acetate and 2.5-3 volumes of cold 100% ethanol. Samples were vortexed and incubated overnight at –80°C. The next day, samples were centrifuged at full speed at 4°C for 30 minutes. The formed pellet was washed twice with 0.5 mL ice-cold 75% ethanol and spun for ten minutes at 4°C. The ethanol was removed, and the pellet was dried and resuspended in 30 mL nuclease-free water.

For each library, 100 to 500 ng/μl of RNA was submitted to the Cornell BRC Genomics Facility for 3’ RNA library preparation and sequencing using Illumina’s NextSeq500, passed through Illumina’s Chastity filter, and files were generated by the Illumina pipeline software v2.18 (Illumina, Inc., San Diego, CA, U.S.A).

### 3’ RNA-Seq analysis

FastQ files were processed using fastp-0.23.2 (cut_right, cut_window_size = 5, cut_mean_quality = 20, length_required = 50) and aligned and counted against the soybean cds sequence, glyma.Wm82.gnm4.ann1.T8TQ.cds.fna.gz collected from SoyBase (Brown *et al.*, 2021), using kallisto v.0.42.4 (bootstrap = 100, mean = 275, sd= 86). Raw read counts were imported to R Studio using tximports, and DESeq was run for genes with summed counts greater than 20 across biological reps. The dataset underwent variance stabilizing transformation for visualization of a PCA plot (Figure 1d).

Pairwise comparisons (alpha = 0.05) were run for each two-whorl combination, whorl-leaf combination, and developmental stage-leaf combination after DESeq analysis. Upregulated and downregulated genes were identified by setting the adjusted p-value (<0.05) and log2fold (>2 and <-2) thresholds. Two-whorl pairwise combination datasets were merged for each whorl to identify genes upregulated in the specific whorl.

### ChIP-Seq sequencing

Mature soybean flowers, buds, and leaves were collected and sent to REquest Genomics (Athens, GA, U.S.A) to immunoprecipitate H3K4 trimethylation DNA, prepare libraries, and sequence a depth of ∼50M reads in a 150 bp format (Ricci *et al.*, 2020).

### ChIP-Seq analysis

Samples were checked for quality using FastQC v.0.12.1 and aligned to the genome glyma.Wm82.gnm4.ann1.T8TQ downloaded from the SoyBase genomic database (Brown *et al.*, 2021) using Bowtie2 v.2.5.1 (Langmead and Salzberg, 2012). Alignment files were converted and sorted using Samtools v.1.18 & Sambamba v.0.7.1 (Li *et al.*, 2009; Tarasov *et al.*, 2015). Peaks were called using MACS2 v.2.2.7.1 (Zhang *et al.*, 2008), and the quality was checked using ChIPQC, R Bioconductor package (Carroll *et al.*, 2014; Gentleman *et al.*, 2004; Zhang *et al.*, 2008). Peak visualization was conducted using KaryotypeR and GenomicFeatures, R Bioconductor packages. Peaks were annotated using GenomicFeatures & ChIPSeeker, R Bioconductor packages (Lawrence *et al.*, 2013; Yu *et al.*, 2015).

## Acknowledgments

This work was made possible by a generous donation from Dr. Phil Needleman, funding from the United Soybean Board (1920-152-0131-D), funding from the Foundation for Food and Agricultural Research New Innovator in Food and Agriculture Research Award (FF-NIA21-0000000017), and funding from the Cornell Graduate School Teaching Assistantship and Research Travel Grant. Soybean plants were tended at Cornell by greenhouse staff, specifically Scott Anthony, and pollinator trials were made possible with the mentorship and support of Dr. Scott McArt and his lab in the Department of Entomology at Cornell University.

## Supporting Data

Supporting Figures: Figures S1-S9

Supporting Tables: Table S1

**Supporting Figure 1:**
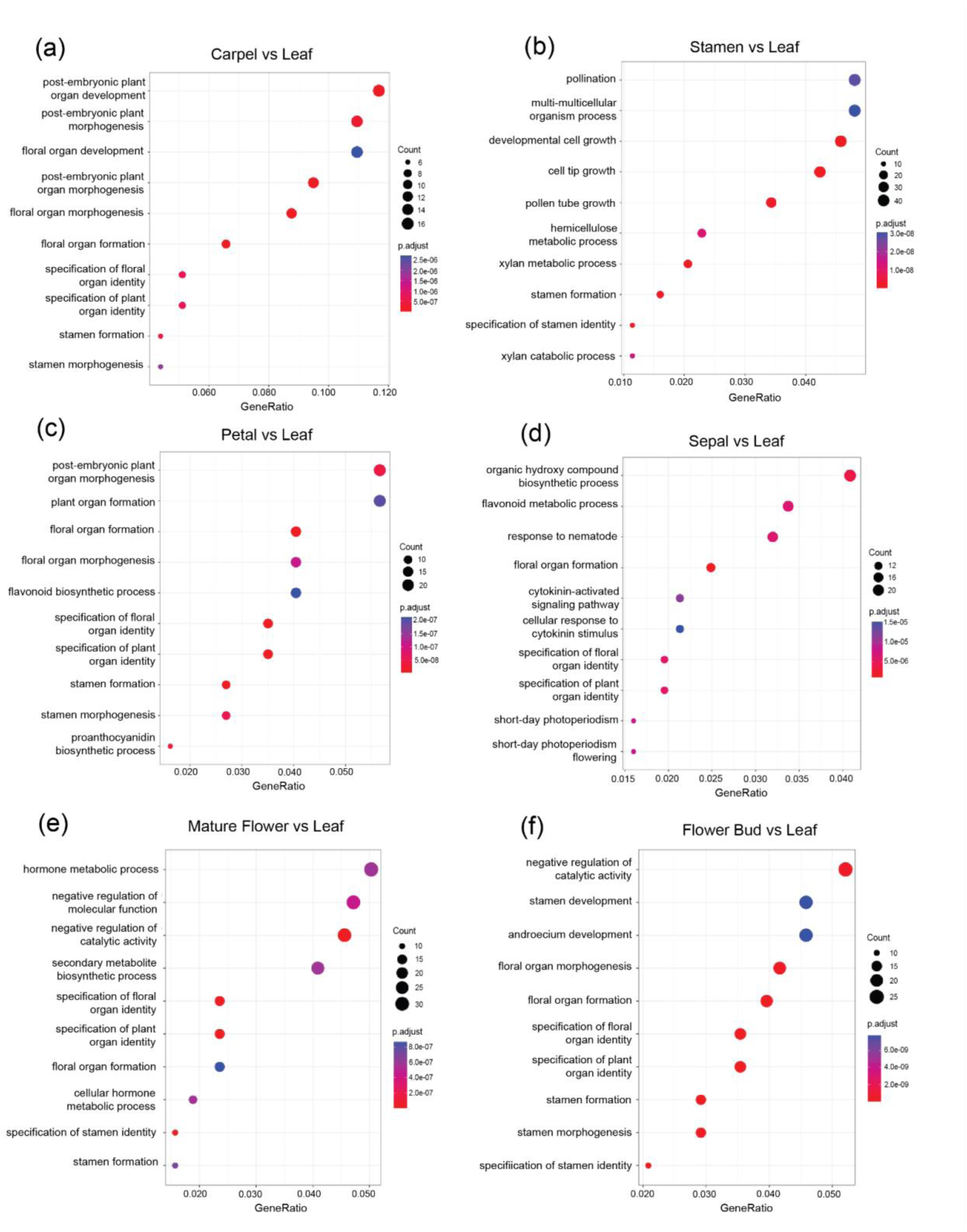
Functions associated with whorl development are highly expressed in RNA-Seq data. GO term analysis lists the most common functions associated with genes that are highly expressed in (a) carpels, (b) stamens, (c) petals, (d) sepals, (e) mature flowers, and (f) flower buds compared to leaves.

**Supporting Figure 2:**
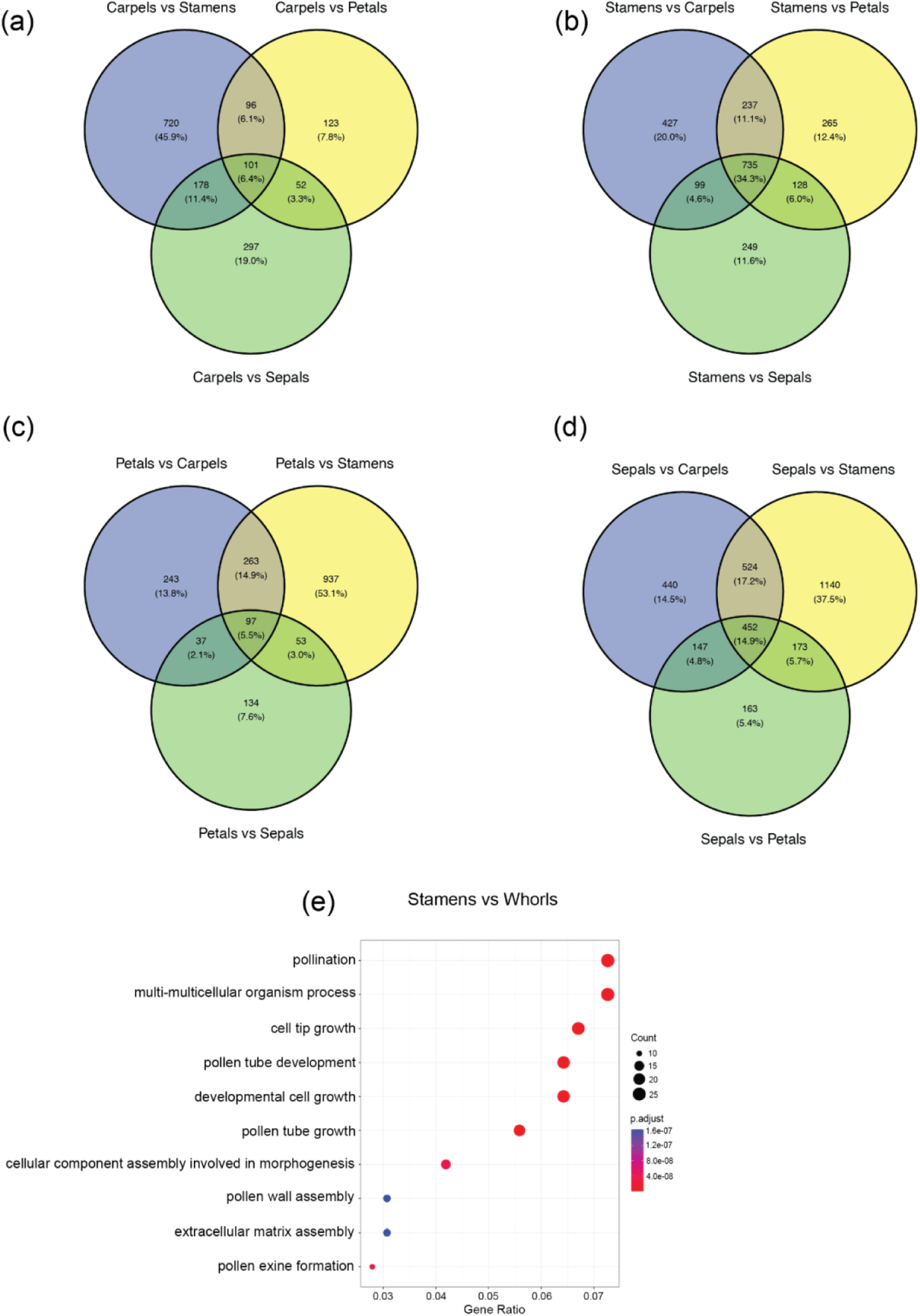
Specific whorl-developmental genes are upregulated only in one whorl. Genes in the center of the Venn diagram are differentially upregulated in only (a) carpels (b) stamens, (c) petals, and (d) sepals. (e) GO term analysis of genes differentially expressed between stamen and the other three whorls.

**Supporting Figure 3:**
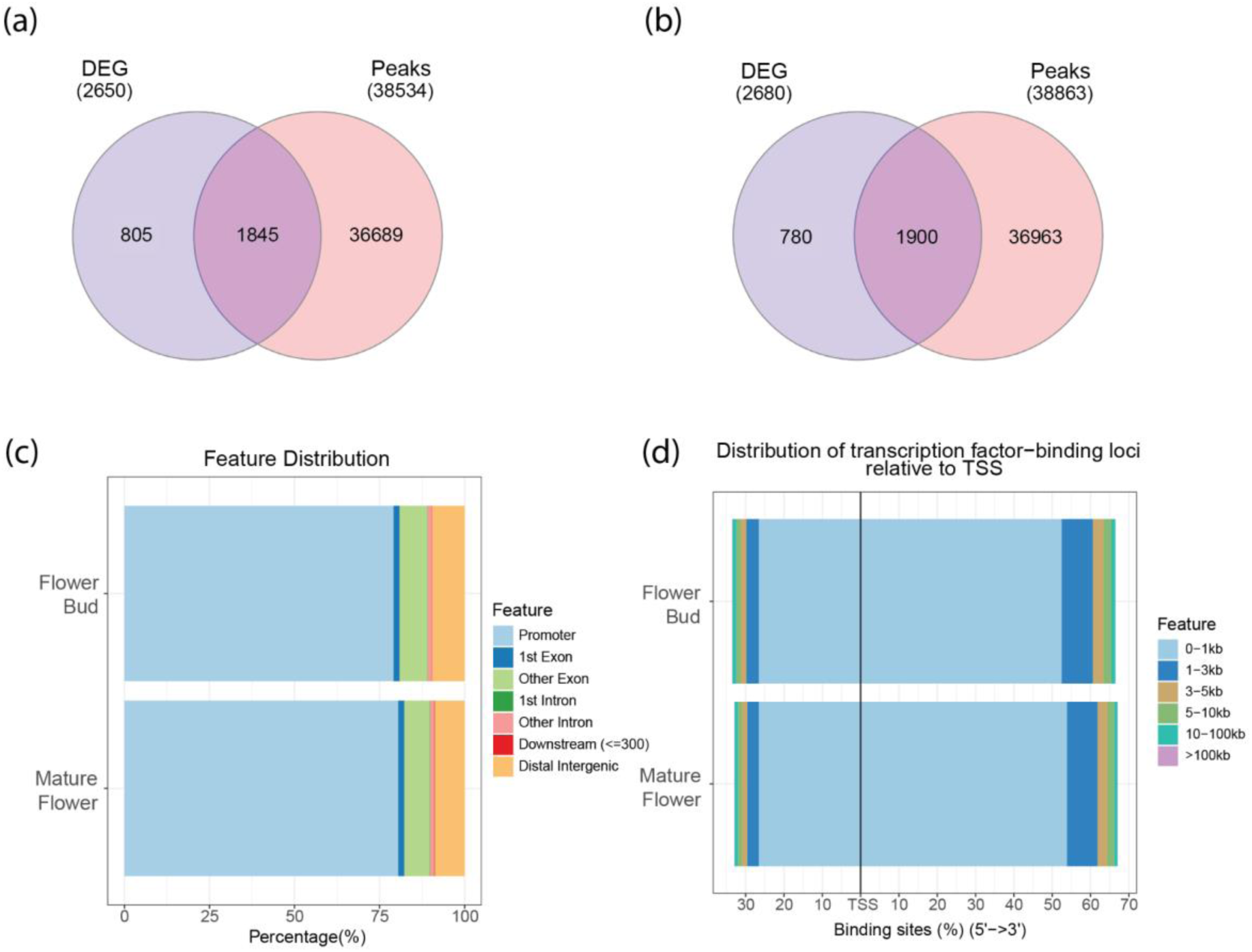
The overlap between differentially expressed genes (DEG) and high quality peaks in flower buds and mature flowers. Venn diagrams between DEG and open chromatin in flower buds (a) mature flowers (b). The gene space feature most abundant in Flower bud and Mature flower is the promoter region (c) and overlap within 1Kb of the transcription start site (TSS) (d).

**Supporting Figure 4:**
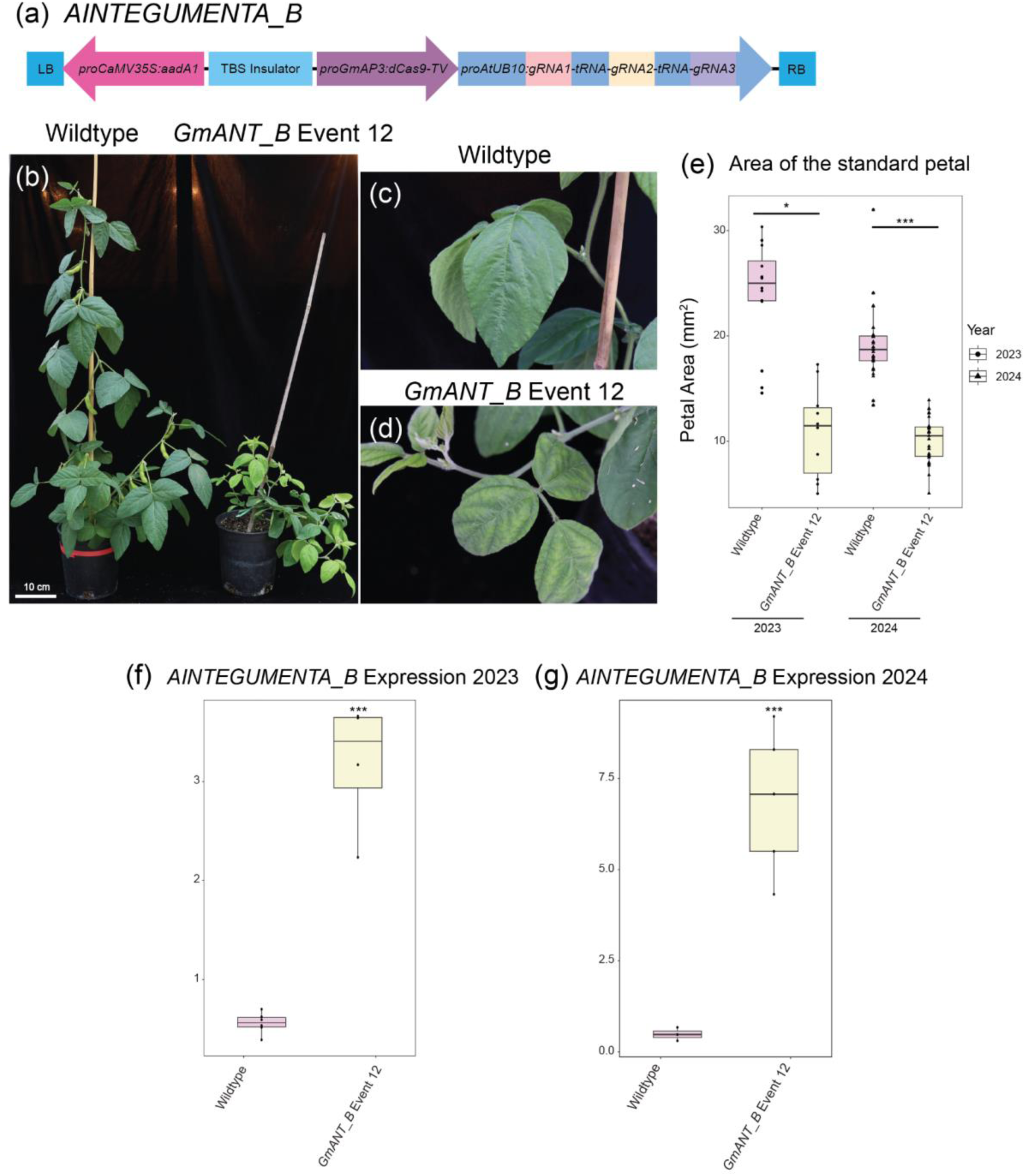
Increased expression of *GmANT B* reduced the width of the standard petal and altered the vegetative phenotype. Construct designs for *GmANT B* (a) using the dCas9-TV system and three gRNAs. (b) Side-by-side comparison of the vegetative phenotype for wildtype and ANT Event 12 plant #28. (c) Image of wildtype leaves. (d) Image of leaves from a plant in ANT Event 12. (e) A boxplot compares the area of wildtype and ANT Event 12 grown in 2023 and 2024. For statistical analysis, samples were fitted to a linear mixed effect model using lmer and the p-value was calculated using a Dunnett’s test. Boxplots measure *ANT_B* expression in 2023 trial (f) and 2024 trial (g). For statistical analysis, samples were fitted to a linear model using ANOVA, and the p-value was calculated with a Dunnett’s test. *** = p < 0.001; ** = p < 0.001; * = p < 0.01

**Supporting Figure 5:**
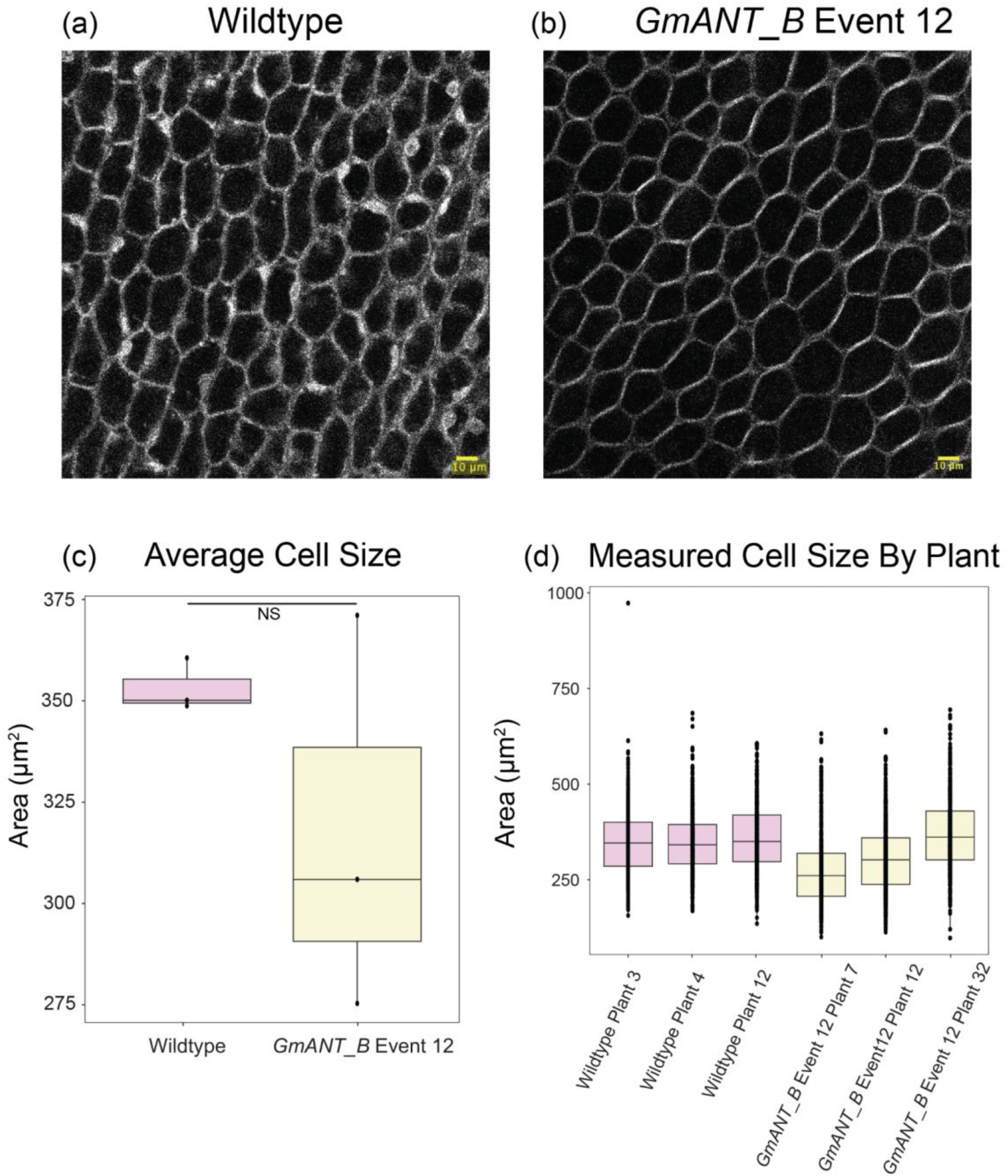
Cell area remains unchanged in *GmANT_B* event 12 petals. 20x magnification confocal microscopy images of (a) wildtype and (b) ANT Event 12 within a 200 micron square scan. (c-d) Cell size was measured using MorpoGraphX software for ∼200 cells per plant and averaged together. Cells partially cut off by the border were not included in the count. For (c), statistical analysis was performed using a Student’s T-Test. (d) Samples were fit to a linear mixed model and non-significant values were calculated with a Tukey test.

**Supporting Figure 6:**
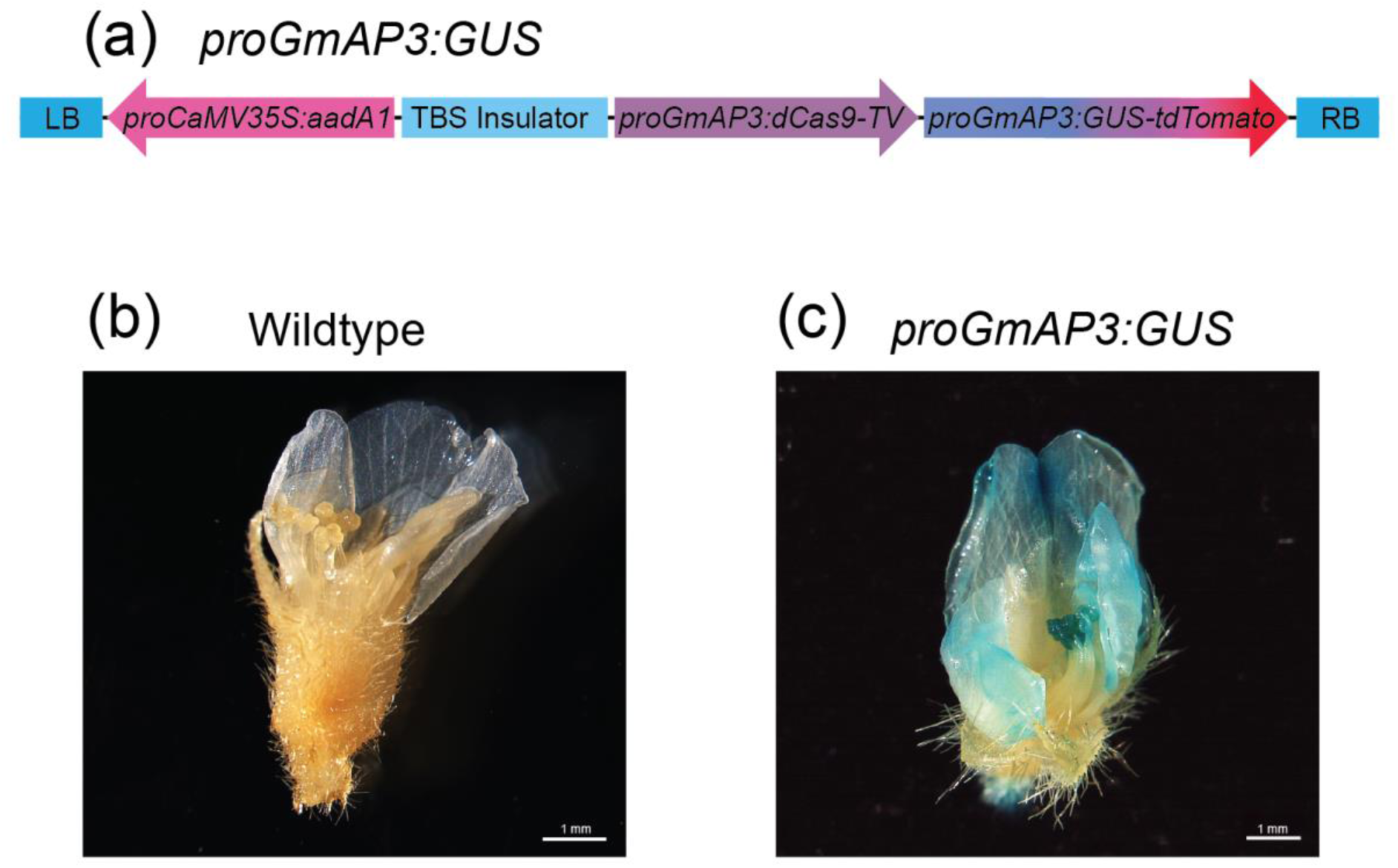
*GmAP3* is expressed in petals and anthers. (a) The construct designed to test the localization of *GmAP3* expression using GUS as a reporter. GUS staining of (a) wildtype and (b) *GmAP3:GUS*.

**Supporting Figure 7:**
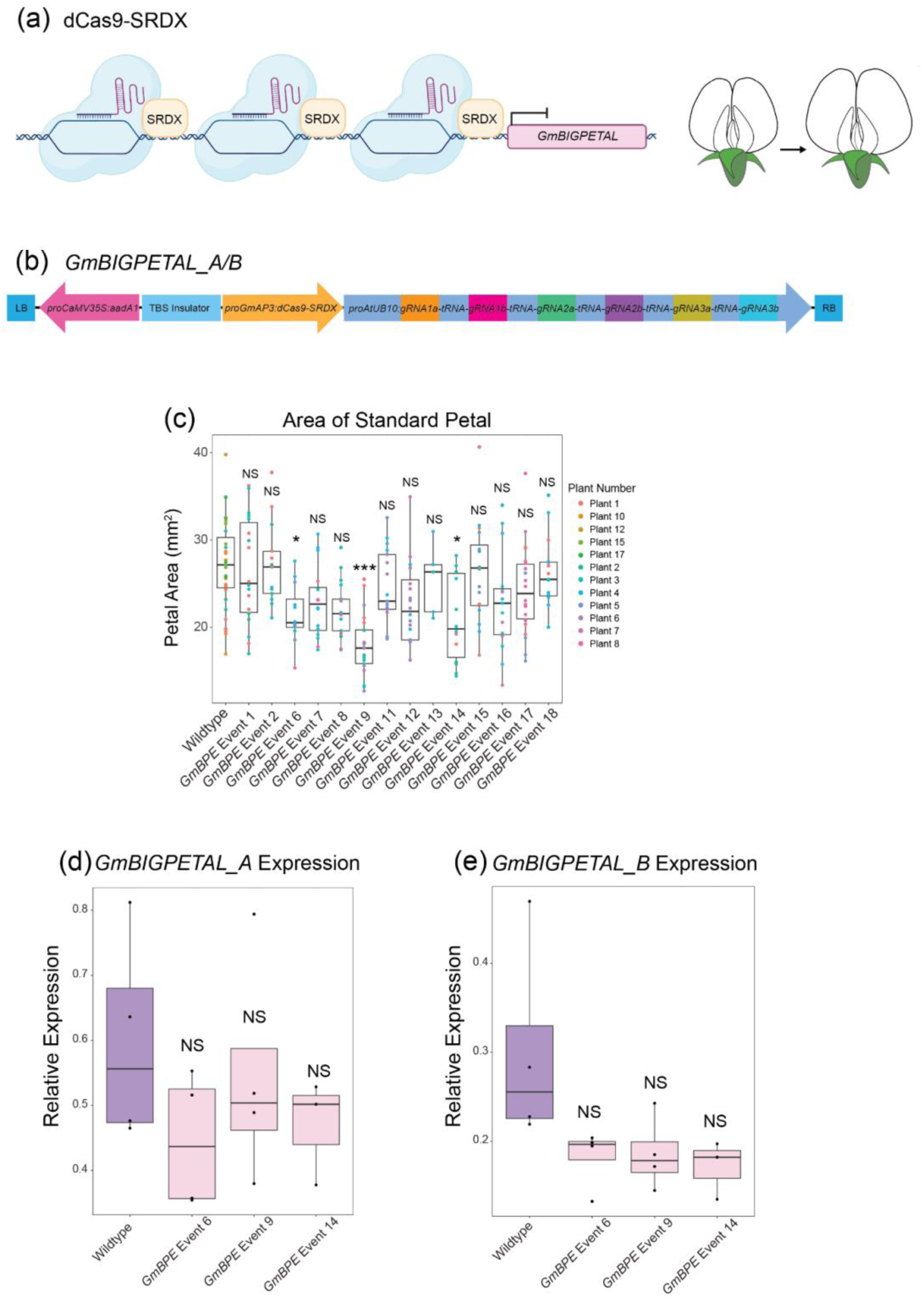
Change in the expression of *BPE* was insignificant in the samples that showed a significant decrease in the petal area. (a) Schematic model of the hypothesis to repress transcription of *BPE* using dCas9-SRDX to increase petal size. (b) The construct is designed to repress the expression of *BPE* using dCas9-SRDX. (c) Boxplots of the area of the standard petal in wildtype and BPE events. For statistical analysis, samples were fitted to a linear mixed effect model using lmer and p-value was calculated using a Dunnett’s test. qRT-PCR expression data for *BPE_A* (d) and *BPE_B* (e) expression in young flower buds. For statistical analysis, samples were fitted to a linear model using ANOVA, and the p-value was calculated with a Dunnett’s test. *** = p < 0.001; ** = p < 0.001; * = p < 0.01

**Supporting Figure 8:**
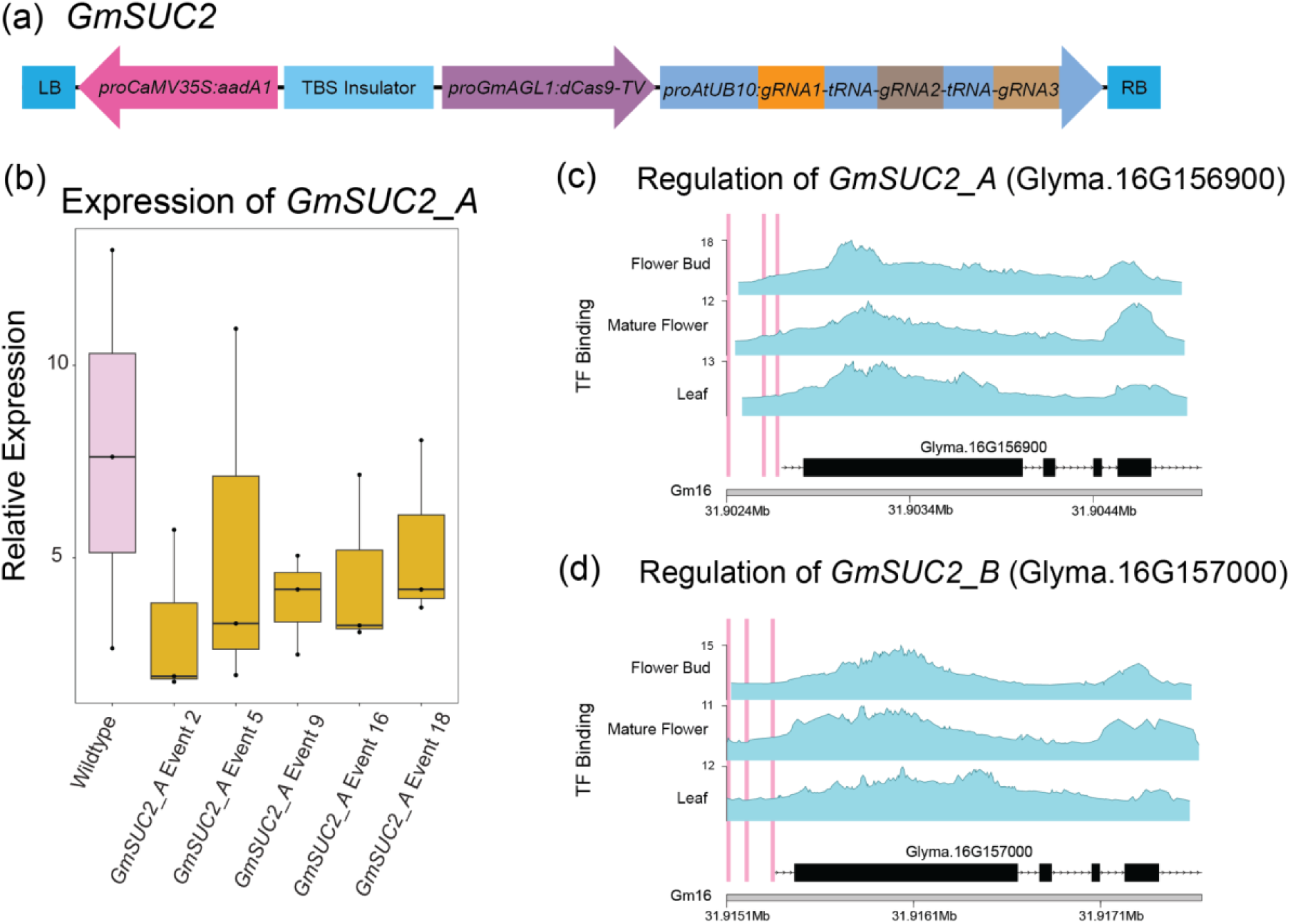
No significant change in expression was observed in plants expressing *GmSUC2_A*. (a) The construct was designed to increase *GmSUC2* expression using the dCas9-TV system and three gRNAs. (b) The change in expression of *GmSUC2_A* quantified using qRT-PCR was not significant. For statistical analysis, samples were fitted to a linear model using ANOVA, and the p-value was calculated with a Dunnett’s test.(c,d) Regions of open chromatin identified by ChIP-Seq in three tissue types (flower bud, mature flower, and leaf) in *GmSUC2_A/B*. The blue peaks represent regions of open chromatin and the pink line indicates the target sites for engineered gRNAs.

**Supporting Figure 9:**
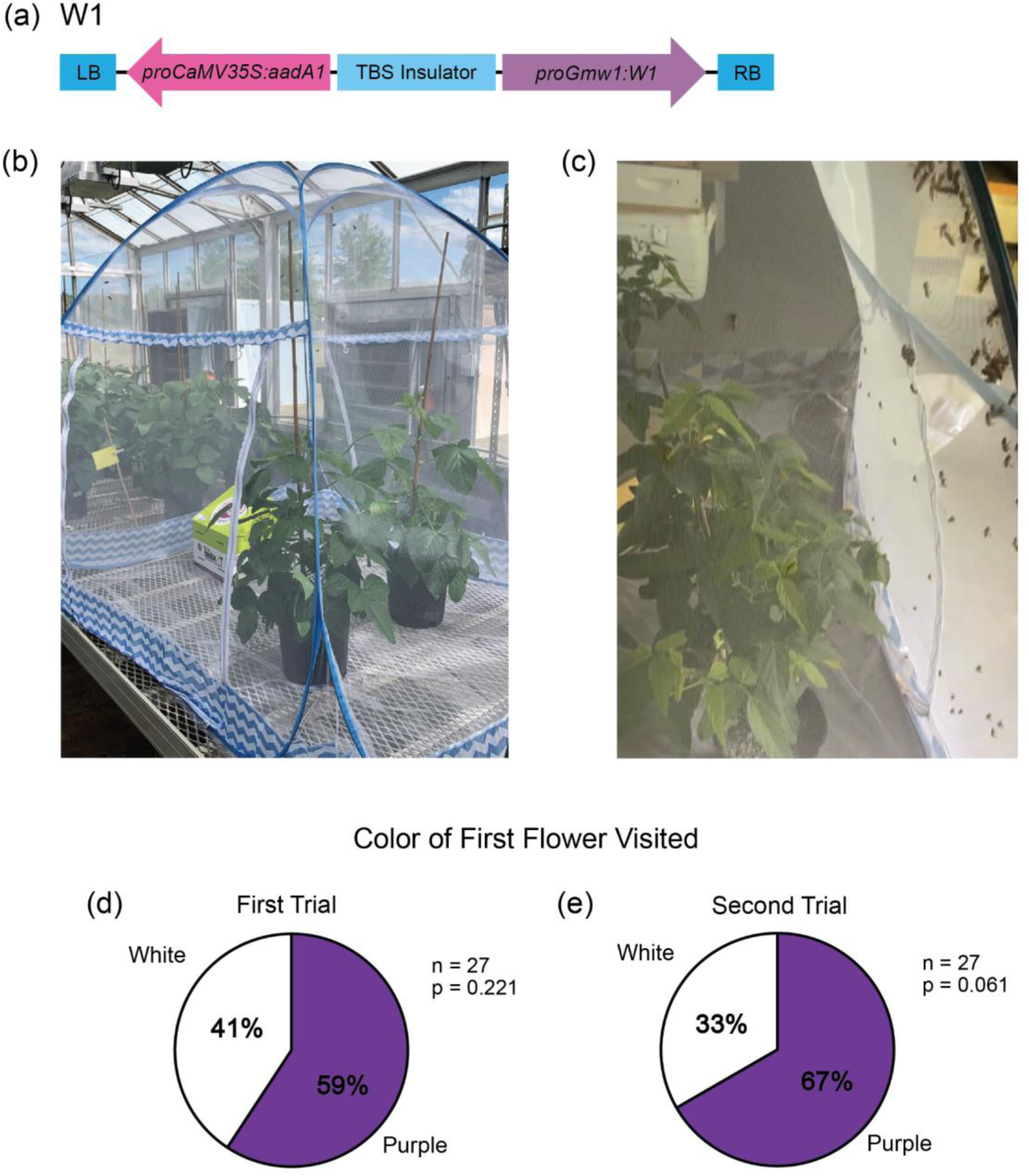
Bees preferred purple flowers. (a) Construct design to produce anthocyanins. (b) Image of plants inside bee cage where the first trial took place with bumblebees. (c) Image of honeybees attempting to escape bee cage. (d,e) Pie charts of the color of the first flower visited for two trials. In the first trial, the door to a hive of *Bombus impatiens* was opened and the color of the first flower-bee interaction was recorded. In the second trial, only one bee was placed in the tent with the two flower choices, and its first flower visit was recorded. Statistics were done using a one-tailed binomial test.

**Supporting Table 1:** Percent sugars in the nectar varied by day and climate. Nectar was extracted between 8:30 and 10:00 AM for Williams 82 (W82) and a control line (WP_1283). To acquire 1.2 ul of nectar, some samples were pooled from flowers across multiple plants. The sugar concentrations were measured using a Brix refractometer.

| Sample Name | Date<br>Extracted | Sucrose<br>Concentration | Light intensity<br>in the<br>greenhouse at<br>8:30 AM | Humidity in<br>the<br>greenhouse at<br>8:30 AM | Temperature in<br>the greenhouse<br>at 8:30 AM |
| --- | --- | --- | --- | --- | --- |
| W82 (Pooled) | 2/13/24 | 12% | 262 uMol | 61.2% | 24.43°C |
| W82 (Pooled) | 2/14/24 | 12% | 348 uMol | 62.1% | 24.71°C |
| WP_1283_2a-1 | 2/15/24 | 8% | 322 uMol | 60.7% | 24.72°C |
| WP_1283_2a-2 | 2/15/24 | 8% | 322 uMol | 60.7% | 24.72°C |
| W82 (Single<br>Plant) | 2/16/24 | 4% | 299 uMol | 64.2% | 22.94°C |
| W82 (Pooled) | 2/16/24 | 5% | 299 uMol | 64.2% | 22.94°C |

